# The M1 and pre-M1 segments contribute differently to ion selectivity in ASICs and ENaCs

**DOI:** 10.1101/2021.02.16.431267

**Authors:** Zeshan P. Sheikh, Matthias Wulf, Søren Friis, Mike Althaus, Timothy Lynagh, Stephan A. Pless

**Author notes:** Correspondence: Stephan A. Pless Department of Drug Design and Pharmacology University of Copenhagen Jagtvej 160, 2100 Copenhagen, Denmark. Department of Natural Sciences, Institute for Functional Gene Analytics, Bonn-Rhein-Sieg University of Applied Sciences, Rheinbach, Germany (Mike Althaus). Sars International Centre for Marine Molecular Biology, University of Bergen, Thormøhlensgate 55, 5006 Bergen, Norway (Timothy Lynagh).

## Abstract

The ability to discriminate between different ionic species, termed ion selectivity, is a key feature of ion channels and forms the basis for their physiological function. Members of the degenerin/epithelial sodium channel (DEG/ENaC) superfamily of trimeric ion channels are typically sodium selective, but to a surprisingly variable degree. While acid-sensing ion channels (ASICs) are weakly sodium selective (sodium:potassium around 10:1), ENaCs show a remarkably high preference for sodium over potassium (>500:1). The most obvious explanation for this discrepancy may be expected to originate from differences in the pore-lining second transmembrane segment (M2). However, these show a relatively high degree of sequence conservation between ASICs and ENaCs and previous functional and structural studies could not unequivocally establish that differences in M2 alone can account for the disparate degrees of ion selectivity. By contrast, surprisingly little is known about the contributions of the first transmembrane segment (M1) and the preceding pre-M1 region. In this study, we use conventional and non-canonical amino acid-based mutagenesis in combination with a variety of electrophysiological approaches to show that the pre-M1 and M1 regions of mASIC1a channels are major determinants of ion selectivity. Mutational investigations of the corresponding regions in hENaC show that they contribute less to ion selectivity, despite affecting ion conductance. In conclusion, our work supports the notion that the remarkably different degrees of sodium selectivity in ASICs and ENaCs are achieved through different mechanisms. The results further highlight how M1 and pre-M1 are likely to differentially affect pore structure in these related channels.

## Introduction

Acid-sensing ion channels (ASICs) and epithelial sodium channels (ENaCs) are members of the degenerin (DEG)/ENaC superfamily of trimeric ion channels and play important roles in neurotransmission and salt homeostasis, respectively. The six human ASIC isoforms known to date (1a, 1b, 2a, 2b, 3 and 4) can form homo- or heterotrimeric proton-gated ion channels and are found primarily in the nervous system, where they contribute to the depolarization of postsynaptic neurons. They are implicated in numerous physiological and pathophysiological processes, including in nociception [1–3], learning and memory [4], fear [5], and neuronal death in ischemic stroke [6, 7]. ENaCs comprise a family of related channels formed by three homologous subunits, α/δ, β and γ, but the canonical channel formed by the α, β, and γ subunits is the most physiologically understood isoform [8]. Canonical ENaCs are located in the apical membrane of epithelial cells, particularly in the kidney, colon and lung, where they contribute to the control of body electrolyte and water homeostasis as well as volume and composition of lung epithelial lining liquid, respectively [9, 10].

Both ASICs and ENaCs are voltage-insensitive channels and share a similar overall subunit topology with intracellular N- and C-termini, two membrane-spanning helices (M1 and M2) and a large extracellular domain [10, 11]. They are inhibited by amiloride and contain sodium-selective pore modules, although the degree of Na^+^/K^+^ permeability greatly varies from *∼*10:1 in ASICs [12] to >500:1 in ENaCs [13, 14]. Functional and structural evidence suggests that in both channel types the ion-conducting pore is lined by M2 [15–18]. Interestingly, this is also the region that displays the highest level of sequence conservation between ASICs and ENaCs. As such, the molecular basis for the pronounced discrepancy in sodium selectivity between the two types of channels remains unclear. In fact, both channel types contain a highly conserved G/SXS motif situated roughly in the middle of the M2, which had been proposed to form a size-exclusion filter [15, 19, 20]. Previous work indeed suggests that the selectivity filter of ENaCs is formed by the S of the G/SXS motif [19] and most ASIC structures available to date display a marked constriction at the level of this highly conserved M2 sequence feature [21, 22]. However, we have recently shown that the ASIC GAS sequence is unlikely to make major contribution to ion selectivity in ASIC1a and 2a, while acidic side chains in the lower M2 appear to be more important [17, 23]. Additionally, the latest ASIC structures show that the lower pore is lined by a reentrant loop formed by side chains from the N-terminus [22], and this pre-M1 region has previously been implicated in ion selectivity in ASICs [24, 25],ENaCs [26, 27] and the related MEC-4/MEC-10 channel [28]. Unlike ASICs, we lack reliable structural information on the pore module of ENaCs, and mutations to conserved acidic side chains in the lower M2 did not have a major impact on ion selectivity [29]. Overall, most previous studies have focused primarily on M2 with regards to contributions to ion selectivity, yet little is known about the contribution of other parts of the pore module, such as M1 and the preceding pre-M1 region. Here, we therefore set out to conduct a comparative analysis of the potential contribution of M1 and pre-M1 to ion selectivity in mouse ASIC1a (m ASIC1a) and human ENaCs (hENaCs).

We employed a combination of site-directed mutagenesis and non-canonical amino acid (ncAA) incorporation in whole cell, single channel and high throughout electrophysiology recordings. Our data suggest that two aromatic residues in ASIC1a M1 play an important role in maintaining a *∼* 10 fold Na^+^/K^+^ permeability ratio. Removal of the bulk of these side chains resulted in a marked decrease in relative Na^+^/K^+^ and Na^+^/Cs^+^ permeability ratios, likely by enlarging the pore diameter. Additionally, single amino acid substitutions of multiple residues in ASIC1a pre-M1 abolish ion selectivity. In contrast, neither the M1 nor the pre-M1 side chains appear to be important for ion selectivity in hENaC, but single channel recordings suggest that some side chains may play a role in ion conduction. Together, our results reveal important yet different roles of the M1 and pre-M1 segment in ASICs and ENaCs, possibly due to differences in their lower pore structures. This consolidates the notion of fundamentally different mechanisms by which these related channel types achieve sodium selectivity.

## Materials and Methods

### Chemicals

NaCl, MgCl_2_, KCl, BaCl_2_, CsCl, HEPES, EGTA, D-glucose, Amiloride hydrochloride hydrate>98%, methylammonium chloride, dimethylamine hydrochloride, ethylammonium chloride, diethylamine hydrochloride, triethylammonium chloride, tetraethylammonium, propylamine hydrochloride and tetrapropylammonium chloride were purchased from SigmaAldrich. Dipropylamine hydrochloride was purchased from Tcichemicals.

### Molecular biology

Mouse ASIC1a (mASIC1a) cDNA, cloned between BamHI and SacI restriction sites of the pSP64 vector, was a gift from Dr. Marcelo Carattino (University of Pittsburgh, US). hENaC *α*, β and γ subunits in the pTNT vector were a gift from Dr. Diego Alvarez de la Rosa (University of La Laguna, Spain). The *α* subunit contained the T334 and A663 polymorphisms. The construct referred to as WT hENaC contains C-terminal truncations in the β and γ subunits (β_R566STOP and γ_K576STOP) to increase expression. Site-directed mutagenesis was performed using custom-designed primers (Eurofins Genomics) and regular PCR with *Pfu*Ultra II Fusion HS DNA Polymerase (Agilent Technologies). Plasmids were linearized using *Eco*RI (for mASIC1a) and *Bam*HI (for hENaC) and used as template for synthesis of mRNA with the Ambion mMESSAGE mMACHINE SP6 transcription kit (Thermo Scientific). ENaCα-mASIC1aM1 chimera DNA cloned between the *Nhe*I and *Xho*l restriction sites in the pUNIV vector was purchased at Twist Bioscience.

### Incorporation of the non-canonical amino acid naphthalene

The nitroveratryloxycarbonyl (NVOC) -protected non-canonical amino acid (ncAA) naphthalene esterified with 5’-O-phosphoryl-2’-deoxycytidylyl-(3’-->5’)adenosine (pdCpA) was incorporated via the nonsense suppression method [30]. Modified *Tetrahymena thermophila* tRNA was prepared by annealing full-length 5’ and 3’ DNA strands (Integrated DNA Technologies, Belgium); RNA was synthetized using the T7-Scribe transcription kit (Cellscript) and purified with Chroma Spin DEPC-H_2_O columns (Clontech, CA, USA). The ncAA naphthalene was ligated to the tRNA with T4 DNA ligase (New England Biolabs, MA, USA). The aminoacylated tRNA was purified with phenol-chloroform extraction and ethanol purification, air dried and store at -80 % until use. The pellet was resuspended in 1 uL water and deprotected with 400 W UV light (Newport UV lamp, # 66921, including Newport power supply # 69920) immediately before injection into oocytes. The deprotected aminoacylated-tRNA was mixed 1:1 with mASIC1a mRNA containing a TAG codon at position 50 and injected into oocytes.

### Oocyte preparation and mRNA injection

Stage IV oocytes were extracted from female *Xenopus laevis* frogs (anaesthetized in 0.3% tricaine, under license 2014-15-0201-00031, approved by the Danish Veterinary and Food administration) by surgery and washed thoroughly in OR-2 medium (in mM: 2.5 NaCl, 2 KCl, 1 MgCl_2_, 5 HEPES, pH 7.4). The oocytes were digested with type I collagenase (1.5 mg/mL, Roche) in OR-2. Stage V and VI oocytes were manually isolated and stored in OR-2 medium. For the injection, oocytes were lined up in a 35 mm dish containing OR-2. For the microinjection, glass capillaries (1.14 mm, 3.5’’, WPI) were pulled using with a horizontal puller (P-1000, Sutter Instruments) and filled with mineral oil (Sigma Aldrich). The tips of the capillaries were broken with fine tweezers to allow RNA uptake. RNA was injected into the oocytes at the oocyte equator (Nanoliter 2010 Injector, World Precision Instruments). RNA was injected in amounts as mentioned in relevant captions and oocytes were incubated in OR-3 (Leibovitz’s L-15 Medium (Gibco) supplemented with 3 mM L-glutamine (Gibco), 0.25 mg/mL gentamycin (Gibco), 15 mM HEPES, pH 7.6) medium for 1-5 days (at 18 C). For ENaC single-channel experiments, ovary lobes were purchased from the European *Xenopus* Resource Centre (Portsmouth, UK) and procedures were approved by the Animal Welfare and Ethical Review Body at Newcastle University (Project ID No: ID 630). RNA-injected oocytes were incubated at 16 °C in a low-sodium solution as previously described [31].

### Electrophysiological experiments and data analysis

For measurements of macroscopic currents through ASICs, oocytes were placed in a custom-built chamber [32], through which bath solution [in mM, 96 NaCl, 2 KCl, 1.8 BaCl_2_, 5 HEPES, pH 7.6 with NaOH) was continuously perfused. For determination of pH concentration-response relationships, the extracellular solution was rapidly and progressively switched to one with lower pH using a gravity-driven ValveBank8 system (AutoMate Scientific, CA, USA). The conditioning pH was 7.6 and after each activating pulse, channels were allowed to recover for 1 min at pH 7.6. Cells were clamped at -20 mV (unless stated otherwise) and currents were recorded with microelectrodes filled with 3 mM KCl (pipette resistance of < 1 MΩ), OC-725C amplifier (Warner, CT, USA) and Digidata 1550 digitizer (Molecular Devices, CA, USA) at 1 kHz with 200 Hz filtering. pH for half-maximal activation (pH_50_) were calculated with the four parameter Hill equation in Prism 6 (GraphPad). For ion selectivity measurements in mASIC1a constructs, IV relationships were determined at a holding potential of -60 mV and applying and a 200-ms voltage ramp from -60 to 60 mV during the peak current. For determination of ion selectivity, reversal potentials with 96 mM extracellular Na^+^, K^+^, Li^+^ and Cs^+^, respectively, were determined using this voltage ramp protocol. Ion permeability ratios were calculated using the measured reversal potentials and a version of the Goldman-Hodgkin-Katz equation (1). pH of the solutions was adjusted using the hydroxide of the respective ions, e.g. NaOH.

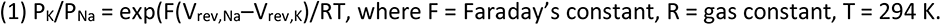

For ion selectivity measurements in hENaC constructs, WT or mutant α, β and γ subunits were injected in a 1:1:1 ratio with a final amount of 5-15 ng mRNA. For construct that showed small macroscopic current sizes, the total RNA amount was increased to 50 ng. Oocytes injected with ENaC constructs were incubated in the presence of 100 μM amiloride to avoid cell death due to excess Na^+^-uptake. For ion selectivity measurements, cells were continuously perfused with bath solution (in mM, 96 XCl, 2 MgCl_2_, 1.8 CaCl_2,_ 5 HEPES; X = Na, K, Li or Cs) containing 100 μM amiloride. Currents were elicited upon switching the solution to one without amiloride. After 1 min., when the currents had reached a steady-state level, a 1 s voltage step protocol was applied ranging from -140 mV to 40 mV. To determine the amiloride-sensitive a current, the protocol was repeated in the presence of 100 μM amiloride. Relative ion permeability ratios were calculated as relative amiloride-sensitive current amplitudes at -100 mV. Data were analyzed in Clampfit 10.7.

Statistical analysis was performed using Graphpad Prism® (7.0) software. An unpaired t-test was used to compare two groups. Multiple groups were compared by one-way ANOVA with a Dunnett’s multiple comparisons test for comparing to a control group (WT). A *p*-value < 0.05 was considered significant. All bars and data points are presented as mean ± standard deviation (SD).

For cell-attached patch-clamp experiments, *Xenopus laevis* oocytes were injected with 10 – 50 ng mRNA. After 24-48 hours, the oocytes were mechanically devitellinized and placed in a recording chamber containing bath solution (in mM: 145 KCl, 1.8 CaCl_2_, 10 HEPES, 2 MgCl_2_ and 5.5 glucose at pH 7.4 adjusted with KOH). Patch pipettes were pulled from borosilicate glass capillaries (6 – 11 MΩ), heat-polished and filled with pipette solution (in mM: 145 NaCl, 1.8 CaCl2, 10 HEPES, 2 MgCl2 and 5.5 glucose). The pH of the pipette solution was adjusted with HCl and NaOH. Current signals were amplified using an LM-PC patch-clamp amplifier (List-Medical, Darmstadt, Germany), low-pass filtered at 100 Hz (Frequency Devices, Haverhill, IL) and recorded at 2 kHz with Axon Clampex software (Axon Instruments, Foster City, CA) using an Axon 1200 interface amplifier. Single-channel analysis was performed with Clampfit version 10.7 (Axon instruments). Recordings were performed at room temperature. Single-channel amplitudes were plotted as a function of voltage, fitted to a linear curve, and values for slope conductances were determined.

Statistical analysis was performed using Graphpad Prism® (7.0) software. To compare two groups an unpaired t-test was used. Multiple groups were compared by one-way ANOVA with a Dunnett’s multiple comparisons test for comparing a control group (WT). a *p*-value < 0.05 was considered significant. All bars and data points are presented as mean ± standard deviation (SD).

### Cell culturing and transfection of HEK293-T for automated patch clamp

ASIC1a K/O HEK293-T cells (provided by Dr. Nina Braun) were grown in monolayer in T75 and T175 flasks (Orange Scientific) in cDMEM (DMEM (Gibco), supplemented with 10% FBS (ThermoFisher) and 1% Penicillin-Streptomycin (ThermoFisher)) and incubated at 37 ⁰C in a humidified 5 % CO_2_ atmosphere. Cells were split at near confluency by treatment with Trypsin-EDTA (ThermoFisher) after washing the cells with PBS. Cells were counted with an EVE™ automatic cell counter (NanoEnTek) and seeded into 10 cm dishes (Orange Scientific) at a density of 2.0 x 10^6^ cells/dish.

For transient transfections in 10 cm dishes, 7 μg of DNA was mixed with 21 μL of Trans-IT LT1 transfection reagent (Mirus) or LipoD transfection reagent and 1750 μL DMEM and incubated at RT for 20 minutes to allow formation of DNA/transfection reagent complexes before addition to the cells. Transfected cells were incubated for 24 hours.

Transfected cells were harvested by Trypsin-EDTA treatment (2 mL/10 cm dish). Detached cells were resuspended in DMEM and centrifuged at 2000 g for 2 minutes. The cells were resuspended in a 1:1 mixture of DMEM and Nanion physiological solution (in mM, 140 NaCl, 4 KCl, 1 MgCl_2,_ 2 CaCl_2,_ 10 HEPES, and 5 Glucose, pH 7.4). Automated patch-clamp experiments were performed 24 hours after transfection.

### Automated patch clamp recordings

The permeability of ASIC1a to a range of monovalent cations (MA^+^: methylammonium^+^, DMA^+^: dimethylamine^+^, EA: ethylammonium^+^, DEA: diethylamine^+^, TEA: trieethylamine^+^, TetraEA^+^: tetraethylamine^+^, PA^+^: propylamine^+^, DPA^+^: dipropylamine^+^, and TetraPA^+^: tetrapropylamine) different in size and shape was assessed with automated whole-cell patch-clamp on the SynchroPatch 384PE. The experiment was designed such that the intracellular solution contained only cations impermeable (or with very low permeability) for ASIC1a, whereas the extracellular solutions contained one of a range of test ions.

Solutions used for the automated patch-clamp experiment are shown below:

| Solution |  | Contents |
| --- | --- | --- |
| Extracellular<br>(ECS) | solution | 140 mM XCl, 4 mM CsCl, 1 mM MgCl<sub>2</sub>, 2 mM BaCl<sub>2</sub>, 10 mM HEPES<br>pH 7.4 and 6, 290 mOsmol/kg |
| Intracellular<br>(ICS) | solution | 80 mM NMDG<sub>2</sub>-SO<sub>4</sub>, 20 mM CsCl, 10 mM HEPES<br>pH 7.2, 270 mOsmol/kg |

The SynchroPatch 394PE consists of a Biomek FX^P^ liquid-handling station (Beckman Coulter) and a planar patch-clamp module (Nanion Technologies), which can be controlled simultaneously via a computer terminal using the PatchControl384 and Biomek software (version 4.1). 25 mL of cells expressing hASIC1a WT or hASIC1a F50A (> 4.0 x 10^6^ cells) were loaded into a Teflon reservoir. Cells were incubated at 20 ⁰C and shaken at 200 rpm in the integrated cell compartment of the SynchroPatch 384PE.

For patch-clamp recordings, 15 μL of cells were loaded into each well of an NPC-384 medium resistance single-hole plate (Nanion Technologies). The plate contains 384 individual wells with a single patch clamp orifice where the cells and ECSs are delivered. The design of the plate and the patch clamp module allows perfusion of the ICS during an experiment. Cells were caught on the patch clamp holes by applying a brief pressure of -100 mBar, and another pulse of pressure (−150 mBar) was applied to reach whole cell configuration. Cells were clamped at 0 mV under atmospheric pressure. Seals of 50 MΩ and above were included in the analysis.

Extracellular solutions (ECS) were provided in 12-well Teflon reservoirs with each column of wells containing an ECS solution with 140 mM of one of the following cations provided as chloride salts shown in Figure 1C. One 12-well dish contained ECS solutions with resting pH (7.4), and another dish contained ECS solutions with activating pH (pH 6). ECS were applied using a liquid stacking approach as described elsewhere [33]. The pipette tips were loaded with 25 μL pH 7.4 solution followed by 15 μL pH 6 solution. For determination of I/V relationships, four sweeps were recorded. For each sweep, the baseline current was recorded for one second (clamped at 0 mV) prior to the application of the activating solution, followed by a delayed dispension with the pH 7.4 solution. This was followed by two washing steps with pH 7.4 solutions after which the protocol was repeated at a different voltage.

**Figure 1:**
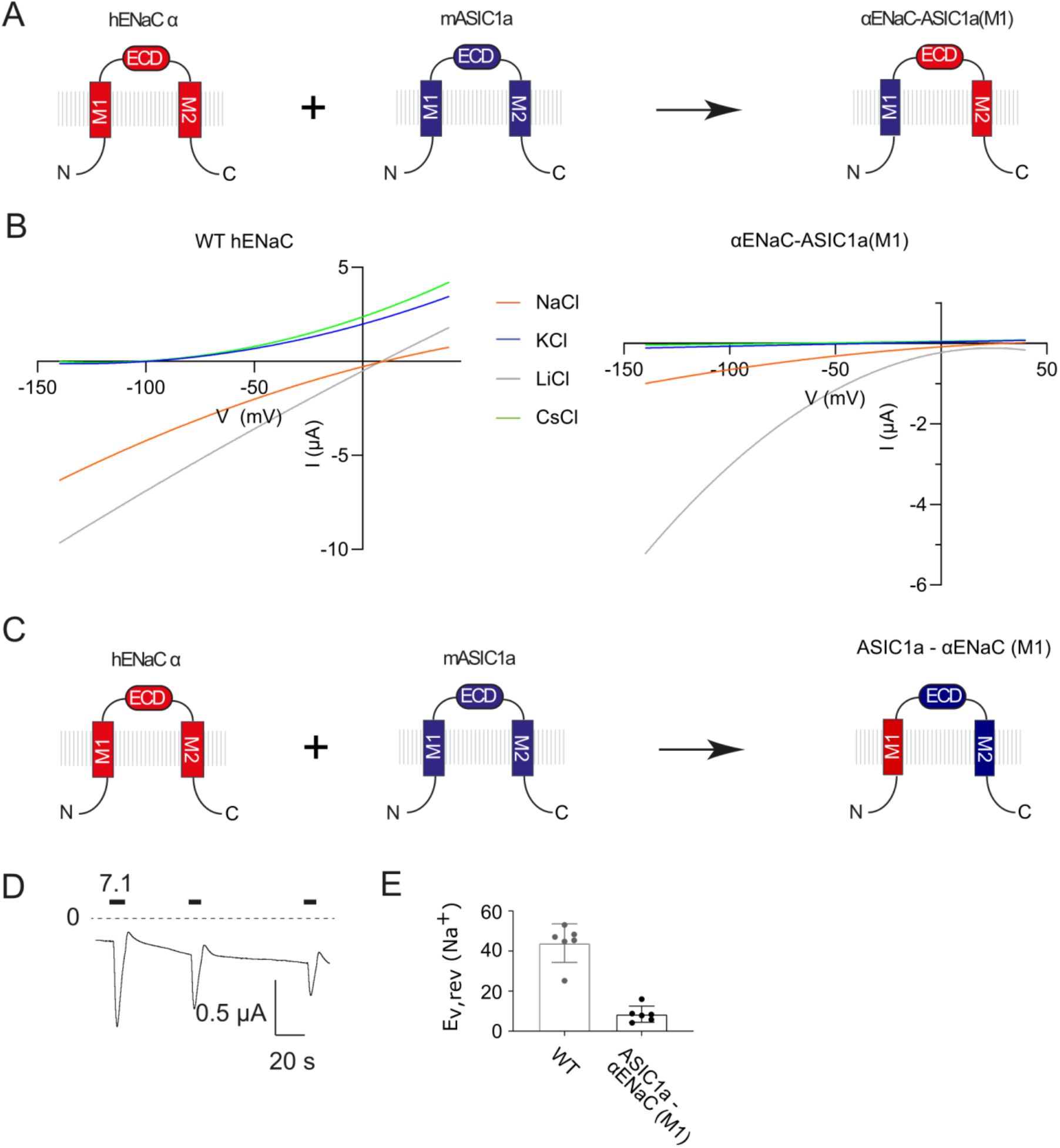
M1 chimeras suggest indirect role of M1 in selectivity. (A) Cartoon illustration of hENaC-mASIC1a chimera design and amiloride-sensitive currents with Na^+^ as the primary extracellular cation. (B) Current-voltage relationships of WT hENaC and hENaC containing the M1 segment of mASIC1a in the α subunit, αENaC-ASIC1a(M1). (C) Cartoon illustration of mASIC1a-hENaC chimera. (D) Current traces of mASIC1a containing the M1 segment of hENaC α, ASIC1a-αENaC(M1). Cells were continuously perfused in ND96 solution (pH 8) and channels were activated with pH 7.1. (E) Reversal potentials with extracellular 96 mM NaCl determined for WT mASIC1a and ASIC1a-αENaC(M1) using a 200 ms voltage ramp from -60 to + 60 mV during the peak current. Currents during the voltage ramps at pH 7.4 were subtracted from currents during activating pH (pH 6).

The Supplementary Material contains Supplementary Figures 1-7.

## Results

### M1-swapped chimeras show altered function

Unlike the highly conserved protein sequences of the pore-lining M2 segments of ASICs and ENaCs, there is noticeable sequence divergence in the M1 segments (Supplementary Figure 1A,B). We hypothesized that differences in the M1 segments between the two channel types may partially account for the observed difference in Na^+^/K^+^ selectivity. To directly test this notion, we swapped the M1 segment of mASIC1a with that of hENaC α subunit, and *vice versa,* (residues A44-F69 in mASIC1a and the equivalent residues in hENaC α) and expressed the resulting chimeras in *Xenopus laevis* oocytes. The αENaC - ASIC1a (M1) chimeric construct, when co-expressed with WT hENaC β and γ, yielded amiloride-sensitive currents (Figure 1A,B) but required injection of 100 ng RNA and three days of incubation for detection of currents. Notably, the αENaC - ASIC1a (M1) construct appeared to be inwardly rectifying. Although the small current magnitudes carried by K^+^ and Cs^+^ prevent us from accurately determining the effects of the M1 swap, the data tentatively suggest an increased permeability for K^+^ and Cs^+^ (I_K+_/I_Na+_ = 0.09 ± 0.06 and I_Cs+_/I_Na+_ = 0.10 ± 0.10 at - 100 mV). By contrast, the Li^+^/Na^+^ amiloride-sensitive current ratio was markedly increased (I_Li_/I_Na_ = 4.2 ± 0.06 compared with 1.2 ± 0.19 in WT hENaC at -100 mV).

The ASIC1a - αENaC (M1) chimera formed constitutively active, pH-sensitive channels that underwent severe tachyphylaxis, rendering an accurate analysis of relative ion permeability ratios of this construct infeasible (Figure 1C-D). However, the measured reversal potential with Na^+^ as the primary cation in the extracellular solution was markedly reduced compared to WT mASIC1a, suggesting that the M1 swap reduces Na^+^ selectivity (Figure 1E).

The altered function of these M1-swapped chimeras suggests that the M1 region contributes to ion selectivity in both channel types. This prompted us to investigate the role of M1 side chains in further detail.

### Residue swapping in the variable region of M1 does not affect ion selectivity of mASIC1a

As pronounced tachyphylaxis precluded obtaining detailed information on relative ion permeabilities of the mASIC1a-hαENaC(M1) chimera, we resorted to swapping non-conserved amino acids in the M1 segment of mASIC1a, one at a time, with those found at corresponding positions in hENaC α (highlighted in red in Figure 2A). We expressed WT or mutant channels in *Xenopus laevis* oocytes and determined ion selectivity by measuring relative ion permeabilities of Na^+^, Li^+^, K^+^ and Cs^+^. Mutations in the more variable region of M1 (residues C3ʹL and G6ʹC-Q20ʹG) had minimal effects on relative ion permeability ratios (Figure 2C, Supplementary Figure 2 and Table 1), suggesting that these residues do not play critical roles in ion selection in mASIC1a.

**Figure 2:**
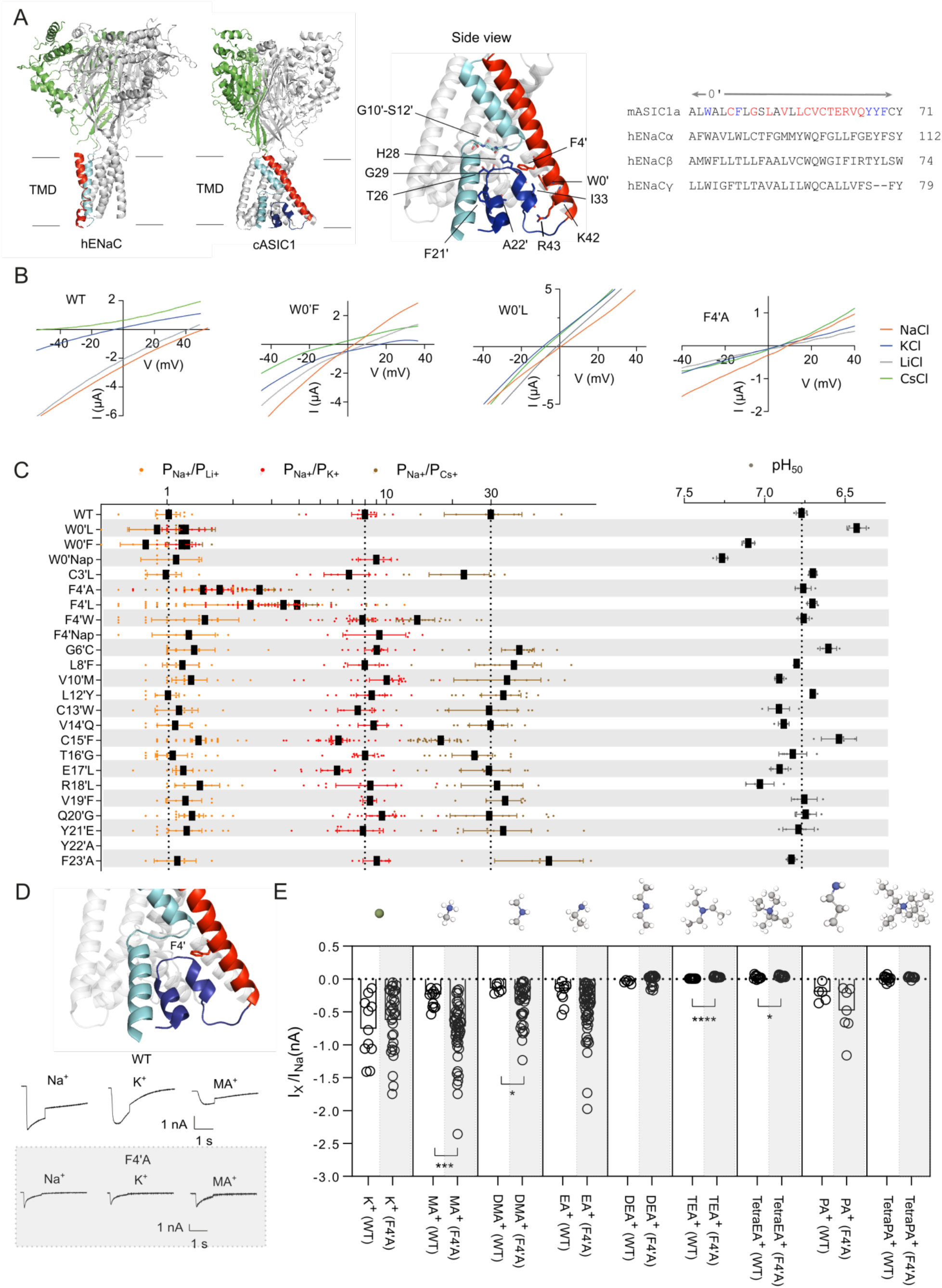
M1 side chains contribute to ion selectivity in mASIC1a. (A) hENaC (PDB: 6BQN) and cASIC1 (PDB: 6VTK) structures (left panel), close-up of cASIC1 structure (middle panel) and sequence alignment of mASI1a and hENaC αβγ M1 (right panel). The residues that were substituted individually for the equivalent residue in hENaC α are shown in red and all aromatics mutated are shown in blue (B) Current-voltage relationships for WT mASIC1a and selected non-selective mASIC1a mutants. (C) Relative ion permeabilities and pH of half-maximum activation (pH_50_) values for mASIC M1 mutants. Data shown as individual data points and mean ± SD. (D) Upper panel shows a close-up view of cASIC1 F4’ (PDB: 6VTK) (M1: red, M2: cyan, pre-M1: dark-purple) and example traces of peak current amplitudes at -20 mV of WT hASIC1a and the non-selective F4’A mutant with different extracellular monovalent cations (lower panel). The solutions containing the difference monovalent cations were applied using a liquid stacking approach, as described elsewhere [33]. (E) Currents measured in HEK293-T cells using a high-throughput patch-clamp robot (SynchroPatch, Nanion). Currents carried by the test ions were compared to those carried by Na^+^ using an unpaired t-test. Note that we cannot exclude the possibility that the absence of permeant ions in the internal solution of SynchroPatch experiments may affect pore structure and cause the lower degree of Na^+^/K^+^ selectivity. Data shown as individual data points and mean ± SD. *, *p* < 0.05; **,*p* < 0.01; ***, *p* < 0.001; ****, *p* < 0.0001. Intracellular solution: in mM; 80 NMDG-SO_4_, 20 CsCl, 10 HEPES, pH 7.2. Extracellular solution: in mM; 140 test ion, 4 CsCl, 1 MgCl_2_, 10 HEPES, pH 7.4 and pH 6, respectively. Abbreviations: MA^+^: methylamine^+^, DMA^+^: dimethylamine^+^, EA^+^: ethylamine^+^, DEA^+^: diethylamine^+^, PA^+^: propylamine^+^.

**Table 1:** Amiloride-sensitive currents with extracellular Na^+^ of hENaC WT and designated mutants. Xenopus laevis oocytes were injected with 5 ng RNA (hENaC WT, α(W4’A)βγ and α(W4’A)β(T4’A)γ or 50 ng RNA (α(W0’F)βγ, α(W0’F)β(W0’F)γ and α(W0’F)β(W0’F)γ(W0’F)) and incubated for 1-3 days. Values presented as mean ± SD; n.d., not determined. Values for the mutants were compared to WT and analyzed using a one-way ANOVA with a Dunnett’s multiple comparisons test. *, *p* < 0.05; **, *p* < 0.01; ***, *p* < 0.001; ****, *p* < 0.0001.

| Construct | I<sub>Na+</sub> (μA) |
| --- | --- |
| hENaC WT | -12.3 ± 4.4 |
| $\alpha(W0'F)\beta\gamma$ | -2.1 ± 0.4 **** |
| $\alpha(W0'F)\beta(W0'F)\gamma$ | -0.7 ± 0.4 **** |
| $\alpha(W0'F)\beta(W0'F)\gamma(W0'F)$ | n.d. |
| $\alpha(W4'A)\beta\gamma$ | -5.7 ± 2.6 ** |
| $\alpha(W4'A)\beta(T4'A)\gamma$ | -3.4 ± 1.7 ** |
| $\alpha(W4'A)\beta(T4'A)\gamma(T4'A)$ | -14.0 ± 5.3 |

### Substitution of aromatic residues in the lower M1 region of mASIC1a affects ion selectivity

Next, we tested the possibility that bulky aromatic side chains in the M1 segments are important in maintaining the structural integrity and/or size of the pore of mASIC1a (highlighted in blue, Figure 2A). We found that substitutions of W0ʹ and F4ʹ altered ion selectivity markedly and decreased functional expression, particularly for the W0’ position (Figure 2C and Tables 1 and 2). Decreasing the side chain volume of W0′ to F or L reduced the Na^+^/K^+^ permeability ratio to unity, caused an alkaline shift in the pH_50_, as well as decreased the rate of desensitization and induced a constitutive current component (Supplementary Figure 2), whereas incorporation of the slightly bulkier non-canonical amino acid (ncAA) naphthalene with the in vivo nonsense suppression method (Supplementary Figure 1C) did not increase ion selectivity but resulted in an alkaline shift in the pH_50_ and increased the rate of desensitization (Supplementary Figure 2). Thus, the bulk of W0′ appears important for both ion selectivity, and gating (Figure 2B/C, Table 2 and Supplementary Figure 2). Substitution of F4′ for smaller side chains (L and A) reduced ion selectivity, but increasing the bulk with a W mutation or incorporation of naphthalene did not increase ion selectivity (Table 2). Contrary to expectations, the relative Na^+^/Cs^+^ permeability ratio was significantly reduced in F4ʹNap compared with WT. This was not investigated further. None of the F4′ mutants affected the pH of half-maximal activation (Figure 2C and Table 2). Hence, the F4′ mutants do not affect gating, but decreasing the bulk of the side chain appears to be detrimental to ion selectivity. Substitution of the aromatic residues in the upper M1 (Y21ʹE-F23ʹA) did not affect ion selectivity or proton sensitivity, although the role of the Y22′A mutant could not be deciphered due to lack of current. In light of the available toxin-stabilized open structure of cASIC1, the lower pore is too wide for the W0’ and F4’ side chains to contribute to ion permeation (Supplementary Figure 5A). However, according to the latest cASIC1 structure in the desensitized state, both W0’ and F4’ are interacting with lower pore-forming residues (Supplementary Figure 5B). In this structure, the absolutely conserved W0’ is making contacts with both a reentrant loop that forms the lower pore of the channel according to this structure, and to residues in the neighboring M2 helix (Supplementary Figure 5B). This complements our data showing that F4’ is important for ion selectivity, whereas the absolutely conserved W0’ is critical for both ion selectivity and gating (Figure 2C and Table 2).

**Table 2:** pH_50_-values and relative ion permeability ratios for mASIC1a M1 substitutions. Values are presented as mean ± SD, and *n* refers to the number of oocytes tested; n.d., not determined. The Y22’A mutant did not produce proton-gated currents. Values for the mutants were compared to WT and analyzed using a one-way ANOVA with a Dunnett’s multiple comparisons test. *, *p* < 0.05; **, *p* < 0.01; ***, *p* < 0.001; ****, *p* < 0.0001.

| Construct | pH<sub>50</sub> | *n* | P<sub>Na+</sub>/P<sub>K+</sub> | P<sub>Na+</sub>/P<sub>Li+</sub> | P<sub>Na+</sub>/P<sub>Cs+</sub> | *n* |
| --- | --- | --- | --- | --- | --- | --- |
| WT | 6.8 ± 0.0 | 5 | 8.0 ± 1.6 | 1.0 ± 0.2 | 29.7 ± 17.2 | 11 |
| W0'Nap | 6.4 ± 0.1**** | 4 | 9.0 ± 1.7 | 1.1 ± 0.3 | n.d. | 5-7 |
| W0'L | 7.1 ± 0.0**** | 4 | 1.2 ± 0.3 **** | 0.9 ± 0.2 | 1.2 ± 0.5 **** | 6 |
| W0'F | 7.3 ± 0.0**** | 5 | 1.2 ± 0.2 **** | 0.8 ± 0.2 | 1.2 ± 0.2 **** | 6 |
| C3'L | 6.7 ± 0.0 | 5 | 6.8 ± 2.0 | 1.0 ± 0.2 | 22.6 ± 8.3 | 8-9 |
| F4'A | 6.8 ± 0.1 | 5 | 1.7 ± 0.6 **** | 1.5 ± 0.7 | 2.6 ± 1.2 **** | 30-33 |
| F4'L | 6.7 ± 0.0 | 5 | 3.4 ± 1.9 **** | 2.4 ± 2.8** | 3.9 ± 2.2 **** | 24-25 |
| F4'W | 6.8 ± 0.0 | 5 | 7.8 ± 2.2 | 1.5 ± 1.5 | 13.8 ± 6.4 ** | 23-24 |
| F4'Nap | n.d. | | 9.3 ± 3.5 | 1.3 ± 0.4 | n.d. | |
| G6'C | 6.6 ± 0.1*** | 5 | 9.0 ± 1.8 | 1.3 ± 0.5 | 40.3 ± 11.4 | 13 |
| L8'F | 6.8 ± 0.0 | 5 | 8.0 ± 1.7 | 1.2 ± 0.3 | 38.1 ± 16.6 | 8-9 |
| V10'M | 6.9 ± 0.0** | 5 | 10.0 ± 3.3 | 1.3 ± 0.4 | 35.6 ± 18.1 | 8-17 |
| L12'Y | 6.7 ± 0.0 | 5 | 8.6 ± 2.1 | 1.0 ± 0.2 | 34.1 ± 10.3 | 11-13 |
| C13'W | 6.9 ± 0.1** | 7 | 7.4 ± 1.8 | 1.1 ± 0.3 | 29.3 ± 13.1 | 9 |
| V14'Q | 6.9 ± 0.0* | 5 | 8.7 ± 1.7 | 1.1 ± 0.2 | 29.8 ± 7.7 | 9 |
| C15'F | 6.5 ± 0.1**** | 4 | 6.0 ± 1.6 | 1.4 ± 0.3 | 17.7 ± 4.8* | 31-34 |
| T16'G | 6.8 ± 0.1 | 5 | 8.0 ± 1.3 | 1.1 ± 0.3 | 25.2 ± 7.9 | 12-13 |
| E17'L | 6.9 ± 0.1** | 5 | 6.0 ± 1.8 | 1.2 ± 0.2 | 29.3 ± 10.4 | 10-16 |
| R18'L | 7.0 ± 0.1**** | 5 | 8.5 ± 3.8 | 1.4 ± 0.4 | 31.9 ± 12.0 | 8-11 |
| V19'F | 6.8 ± 0.1 | 4 | 8.4 ± 0.7 | 1.2 ± 0.3 | 34.8 ± 7.6 | 8 |
| Q20'G | 6.7 ± 0.1 | 5 | 9.5 ± 2.8 | 1.3 ± 0.3 | 29.4 ± 17.1 | 14-16 |
| Y21'E | 6.8 ± 0.1 | 5 | 7.8 ± 3.2 | 1.2 ± 0.4 | 34.1 ± 18.4 | 12-14 |
| Y22'A | n.d. |  | n.d. | n.d. | n.d. |  |
| F23'A | 6.8 ± 0.0 | 5 | 9.0 ± 1.3 | 1.1 ± 0.3 | 55.0 ± 22.8 | 6-7 |

### Permeability properties of WT ASIC1a and the non-selective F4**′**A mutant

To test the hypothesis that the removal of side chain bulk of aromatic residues in the lower ASIC1a M1 affects the size of the pore, we set out to measure currents carried by a range of monovalent cations with different sizes through WT and the non-selective F4’A mutant. We chose the F4ʹA mutant based on the fact that it produced larger currents in X*enopus laevis* oocytes and because it does not obviously affect channel gating (unlike e.g. W0ʹF and W0ʹL). HEK-293T cells without endogenous hASIC1a channels [34] were transfected with WT or F4ʹA hASIC1a DNA and currents were recorded using an automated patch-clamp system (SyncroPatch 384PE). Similar to conventional patch-clamp, the SyncroPatch 384PE system allows control over both the intracellular and extracellular solutions, but with greater overall throughput. We used *N*-methyl-D-glucamine (NMDG^+^) as the only cation in the intracellular solution to ensure minimal outward ionic flow from the intracellular solution and thus facilitate isolation of small inward currents carried by the extracellular test ions. pH 6-induced currents were recorded at four test potentials, -20, 10, 40 and 70 mV, to estimate the reversal potentials with different extracellular monovalent test ions (Supplementary Figure 3). Figure 2E shows the pH 6-induced inward currents at -20 mV carried by the different monovalent test ions normalized to the current carried by Na^+^. Based on these data, the F4′A mutant conducts larger monovalent cations more readily compared to WT hASIC1a (Figure 1E and Table 3). For example, the current carried by methylammonium^+^ (MA^+^) in F4′A is almost equal in magnitude to the current carried by Na^+^ (I_MA_/I_Na_ = 0.81 ± 0.45, p = 0.29), whereas in WT, the magnitude of the MA^+^ current is only one third of the Na^+^ current (I_MA+_/I_Na+_ = 0.31 ± 0.14; p < 0.001) (Figure 2D/E). Also, the permeability of dimethylamine (DMA^+^) is significantly increased in F4ʹA (I_DMA+_/I_Na+_ = 0.40 ± 0.31) compared with WT (I_DMA+_/I_Na+_ = 0.14 ± 0.07; p < 0.05). A similar pattern was observed when we examined the reversal potentials with the respective extracellular test ions. Supplementary Figure 3C shows reversal potentials estimated with different monovalent cations. The corresponding relative ion permeability ratios are reduced in the F4ʹA mutant for all ions, except for EA^+^ (Supplementary Figure 3D). This suggests that the pore of the F4′A mutant is larger than that of the WT channel.

**Table 3:**
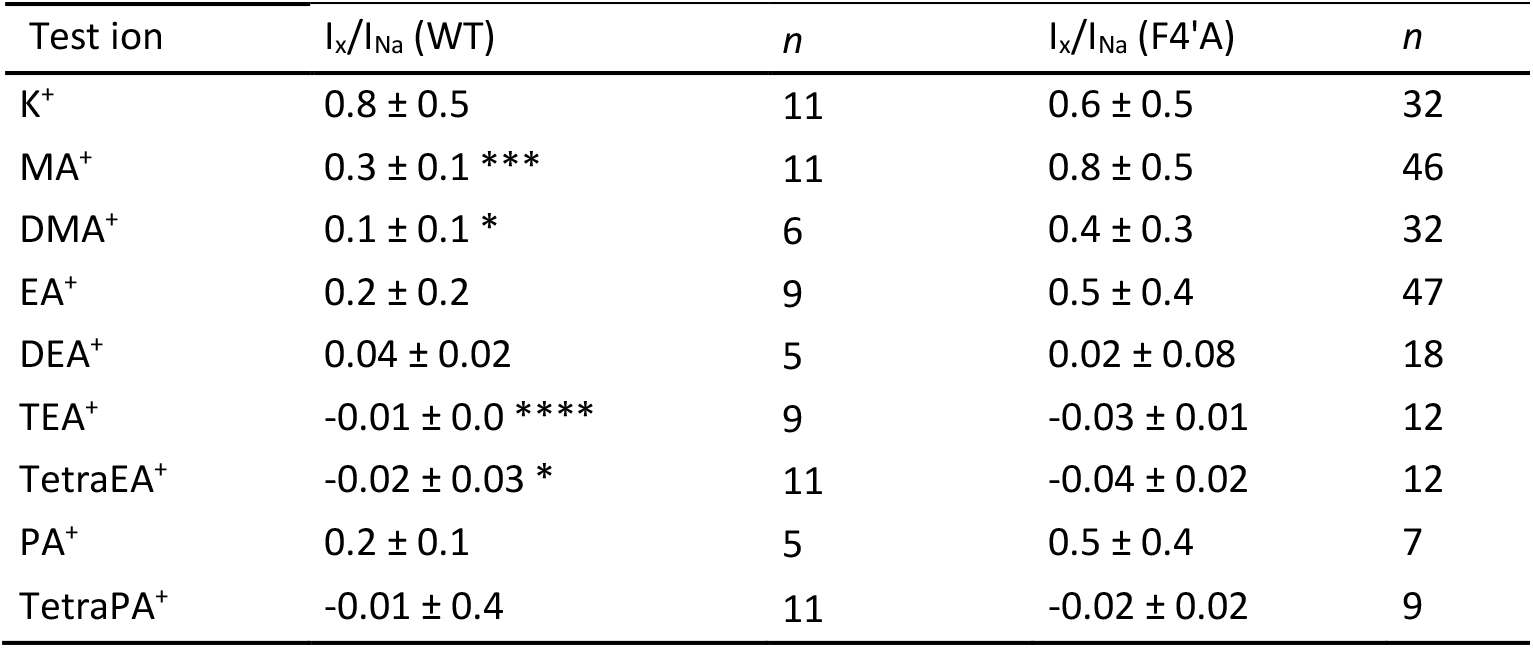
Normalized pH 6-induced inward peak currents carried by extracellular monovalent cations in hASIC1a WT and F4’A. Currents were recorded on a high-throughput patch clamp robot (SynchroPatch 384PE). Currents carried by the different test ions were normalized to the mean Na^+^ current. Values presented as mean ± SD. Currents carried by the test ions were compared to those carried by Na^+^ using an unpaired t-test. *, *p* < 0.05; **, *p* < 0.01; ***, *p* < 0.001; ****, *p* < 0.0001.

Our mASIC1a data shows that substitution of the F4′ side chain significantly affects the size of the pore and increases the permeability towards larger monovalent cations. This finding supports the notion that F4′, and possibly W0′, can indirectly affect ion selectivity. As aromatic residues at the 0′ positions are conserved in all three hENaC subunits and the 4′ residue in hENaC α is also an aromatic, we hypothesized that these residues in hENaC might play an equally important role in ion selectivity.

### The 0ʹ and 4ʹ residues of hENaC M1 are unlikely to be critical for ion selectivity

Based on the mutational screen performed on the M1 segment of mASIC1a, we concluded that side chains at the 0′ and 4′ positions are critical for ion selectivity. We therefore decided to investigate the role of the corresponding residues in hENaC. hENaC α-, β- and γ subunits were expressed in *Xenopus laevis* oocytes, and we measured currents using two-electrode voltage clamp. The voltage protocol and I/V relationships for WT hENaC and a designated mutant in either 96 mM NaCl, KCl, LiCl or CsCl in the bath solution are shown in Figure 3A-C. For WT hENaC, we found that Na^+^ and Li^+^ currents reverse around 25 mV, whereas Cs^+^ and K^+^ are virtually non-permeable, as the Cs^+^ and K^+^ currents reverse at V_m_ < -100 mV. This is consistent with previously observed values for Na^+^/K^+^ and Na^+^/Cs^+^ current ratios [14, 35]. As reversal potentials could not be accurately determined, relative ion conductance ratios for all hENaC constructs were estimated as amiloride-sensitive K^+^, Li^+^ and Cs^+^ currents measured at -100 mV normalized to the amiloride-sensitive Na^+^ current [19, 36].

**Figure 3:**
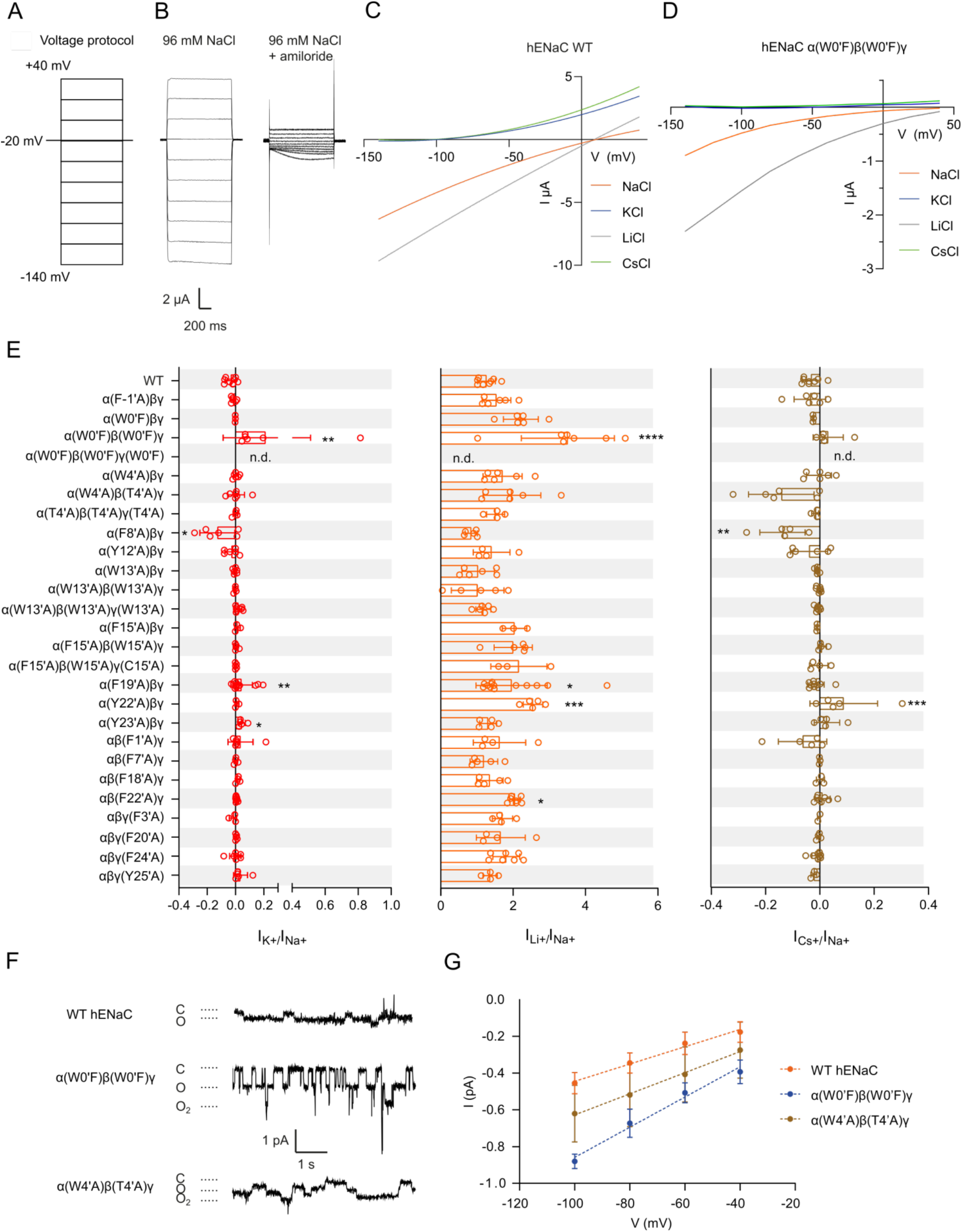
M1 mutations in hENaC impact conductance, but have only minimal effects on selectivity. (A) Voltage protocol for relative amiloride-sensitive current measurements with different extracellular cations. *Xenopus laevis* oocytes were injected with hENaC WT or mutant cRNA and macroscopic currents were recording using two-electrode voltage clamp. *Xenopus laevis* oocytes were clamped at -20 mV and continuously perfused with solution containing amiloride (100 μM), and currents were elicited upon switching the extracellular solution to one without amiloride. After 1 min., the voltage-protocol was applied. Currents were measured during 1 s voltage steps from a holding potential of -20 mV to test potentials of -140 mV to +40 mV in 20 mV increments. Currents measured in the presence of 100 µM amiloride (C) were subtracted from currents measured in the absence of amiloride (B). (C-D) Current-voltage relationships of the amiloride-sensitive current in 96 mM Na^+^, K^+^, Li^+^ and Cs^+^ bath of WT hENaC and the designated mutant. (E) Relative I_K+_/_Na+_ (red), I_Li+_/I_Na+_ (orange) and I_Cs+_/I_Na+_ (brown) current amplitudes measured at –100 mV in WT hENaC and the designated mutants. Values shown as mean ± SD. Data were compared by a one-way ANOVA with a Dunnett’s multiple comparisons test for comparison with control (WT hENaC). *, *p* < 0.05; **, *p* < 0.01; ***, *p* < 0.001. (F) Single-channel traces of WT hENaC and designated mutants in the cell-attached configuration in *Xenopus laevis* oocytes recorded at -100 mV. The spikes originate from endogenous mechanosensitive ion channels. (G) Single-channel I/V relationships for the channels shown in (F). Slopes estimated by linear regression gave conductances of: WT = 4.7 pS ± 0.98; α(W0’F)β(W0’F)γ = 8.6 pS ± 1.0; (αW4’A)(βT4’A)γ = 5.3 pS ± 2.2. Data presented as mean ± SD for 4 to 7 patches. The value for α(W0’F)β(W0’F)γ is significantly greater than that of the WT channel (*p*-value = 0.0004).

Single amino acid substitutions were introduced at the 0′ and 4′ positions in one, two or all three hENaC subunits. W0′L or W0′A substitutions in hENaC α co-expressed with WT β and WT γ rendered the channel non-functional, but the W0′F mutation in hENaC α expressed with WT β and WT γ yielded amiloride-sensitive currents and showed relative amiloride-sensitive currents similar to WT hENaC (Figure 3E and Table 5). W0′F substitutions in both α and βENaC (expressed with WT γ) increased the amiloride-sensitive Li^+^/Na^+^, but had no significant effect on the K^+^/Na^+^ or Cs^+^/Na^+^ currents (Figure 3E and Table 5). With W0′F substitutions in all three hENaC subunits, amiloride-sensitive currents could not be determined with any of the four ions due to lack of current. Although we cannot comment on the triple mutant scenario, W0′ does not appear to contribute to ion selectivity in hENaC, at least upon W0′ substitutions in one or two out of three subunits. Substitution of the amino acids at the 4′ position in α-, β- and γENaC yielded measurable amiloride-sensitive currents for all three mutants, but the relative amiloride-sensitive current profiles of all tested 4′ position mutations are similar to that of WT hENaC (Figure 3E and Table 5).

Although mutations at the 0′ and 4′ positions of hENaC had no significant effect on ion selectivity, there is a possibility that other amino acids in M1 of hENaC contribute to ion selectivity. The M1 of hENaC contains a higher number of aromatic residues compared with mASIC1a (Figure 1A). We set out to test the hypothesis that the larger number of aromatic residues explains why hENaC is more Na^+^ selective, possibly by indirectly decreasing the pore diameter. To this end, single amino acid substitutions were introduced individually into the M1 of hENaC. For positions that contained an aromatic amino acid in two or three subunits, substitutions were made in two or all three subunits. Figure 3E depicts amiloride-sensitive currents with Li^+^, K^+^ or Cs^+^ relative to the current carried by Na^+^. None of the mutants exhibited significantly altered ion selectivity profiles and all remained non-permeable to K^+^ and Cs^+^, except for α(F19ʹA)βγ and α(Y23ʹA)βγ, which showed a slight permeability for K^+^ and Cs^+^ (Figure 3E and Table 5). Some mutants exhibited modestly altered Li^+^/Na^+^ current. These include α(F19ʹA)βγ, α(Y22ʹA)βγ αβ(F22ʹA)γ (Figure 3E and Table 4).

**Table 4:** Single channel conductances of WT hENaC, 0’ and 4’ mutants. WT and mutant hENaC RNA was injected in *Xenopus laevis* oocytes and single-channel conductances were determined in the cell-attached patch clamp configuration with 140 mM extracellular Na^+^. Single channel conductances presented as mean ± SD and were compared to WT ENaC using a one-way ANOVA with a Dunnett’s multiple comparisons test. *, *p* < 0.05; **, *p* < 0.01; ***, *p* < 0.001; ****, *p* < 0.0001

| Construct | Single channel conductance (pS) | *n* |
| --- | --- | --- |
| WT hENaC | 4.7 ± 1.0 | 6 |
| $\alpha$(W0'F)$\beta\gamma$ | 7.4 ± 3.1 * | 5 |
| $\alpha$(W0'F)$\beta$(W0'F)$\gamma$ | 8.6 ± 1.0 *** | 6 |
| $\alpha$(W4'A)$\beta\gamma$ | 6.6 ± 1.0 | 4 |
| $\alpha$(W4'A)$\beta$(T4'A)$\gamma$ | 5.3 ± 2.2 | 7 |

**Table 5:** Relative amiloride-sensitive currents in WT hENaC and hENaC M1 mutants. Relative amiloride-sensitive currents determined for with WT ENaC and hENaC mutants containing the designated amino acid substitutions at positions with aromatic side chain. Data shown as mean ± SD;; n.d., not determined and *n* refers to the number of individual oocytes. Values for mutants were compared to WT hENaC using a one-way ANOVA with a Dunnett’s multiple comparisons test. *, *p* < 0.05; **, *p* < 0.01; ***, *p* < 0.001; ****, *p* < 0.0001.

| Construct | $I_K/I_{Na}$<br>(-100mV) | $I_{Li}/I_{Na}$<br>(-100mV) | $I_{CS}/I_{Na}$<br>(-100mV) | $n$ |
| --- | --- | --- | --- | --- |
| WT hENaC | $-0.04 \pm 0.04$ | $1.2 \pm 0.2$ | $-0.04 \pm 0.04$ | 7 |
| $\alpha(W-1'A)\beta\gamma$ | $-0.02 \pm 0.02$ | $1.5 \pm 0.4$ | $-0.04 \pm 0.06$ | 6 |
| $\alpha(W0'F)\beta\gamma$ | $-0.02 \pm 0.00$ | $2.2 \pm 0.5$ | $-0.03 \pm 0.00$ | 6 |
| $\alpha(W0'F)\beta(W0'F)\gamma$ | $0.2 \pm 0.3$ ** | $3.5 \pm 1.3$ **** | $0.03 \pm 0.06$ | 5 |
| $\alpha(W0'F)\beta(W0'F)\gamma(W0'F)$ | n.d. | n.d. | n.d. | |
| $\alpha(W4'A)\beta\gamma$ | $0.00 \pm 0.02$ | $1.7 \pm 0.5$ | $0.0 \pm 0.1$ | 6 |
| $\alpha(W4'A)\beta(T4'A)\gamma$ | $0.00 \pm 0.06$ | $2.0 \pm 0.9$ | $-0.1 \pm 0.1$ * | 3-4 |
| $\alpha(W4'A)\beta(T4'A)\gamma(T4'A)$ | $0.00 \pm 0.02$ | $1.5 \pm 0.3$ | $-0.02 \pm 0.01$ | 4 |
| $\alpha(F8'A)\beta\gamma$ | $-0.1 \pm 0.1$ * | $0.9 \pm 0.2$ | $-0.1 \pm 0.1$ ** | 5-6 |
| $\alpha(Y12'A)\beta\gamma$ | $-0.03 \pm 0.05$ | $1.4 \pm 0.5$ | $-0.04 \pm 0.1$ | 4-6 |
| $\alpha(W13'A)\beta\gamma$ | $-0.00 \pm 0.01$ | $1.0 \pm 0.5$ | $-0.01 \pm 0.01$ | 6 |
| $\alpha(W13'A)\beta(W13'A)\gamma$ | $-0.00 \pm 0.01$ | $1.0 \pm 0.7$ | $-0.00 \pm 0.01$ | 4-7 |
| $\alpha(W13'A)\beta(W13'A)\gamma(W13'A)$ | $0.02 \pm 0.02$ | $1.2 \pm 0.2$ | $-0.01 \pm 0.01$ | 6-8 |
| $\alpha(F15'A)\beta\gamma$ | $0.01 \pm 0.02$ | $2.0 \pm 0.3$ | $-0.01 \pm 0.00$ | 3 |
| $\alpha(F15'A)\beta(W15'A)\gamma$ | $0.01 \pm 0.01$ | $2.0 \pm 0.5$ | $0.01 \pm 0.02$ | 4-5 |
| $\alpha(F15'A)\beta(W15'A)\gamma(C13'A)$ | $0.00 \pm 0.01$ | $2.2 \pm 0.8$ | $-0.03 \pm 0.01$ | 3 |
| $\alpha(F19'A)\beta\gamma$ | $0.04 \pm 0.08$ ** | $2.0 \pm 1.0$ * | $-0.01 \pm 0.03$ | 9-12 |
| $\alpha(Y22'A)\beta\gamma$ | $0.01 \pm 0.01$ | $2.6 \pm 0.3$ *** | $0.09 \pm 0.1$ *** | 3-5 |
| $\alpha(Y23'A)\beta\gamma$ | $0.05 \pm 0.03$ * | $1.3 \pm 0.2$ | $0.03 \pm 0.05$ | 5 |
| $\alpha\beta(F1'A)\gamma$ | $0.0 \pm 0.0$ | $1.3 \pm 0.2$ | $-0.03 \pm 0.04$ | 3-4 |
| $\alpha\beta(F7'A)\gamma$ | $0.0 \pm 0.01$ | $1.2 \pm 0.4$ | $-0.00 \pm 0.00$ | 3-5 |
| $\alpha\beta(F18'A)\gamma$ | $0.02 \pm 0.01$ | $1.4 \pm 0.4$ | $0.00 \pm 0.02$ | 3-5 |
| $\alpha\beta(F22'A)\gamma$ | $0.01 \pm 0.01$ | $2.1 \pm 0.2$ * | $0.01 \pm 0.03$ | 7-8 |
| $\alpha\beta\gamma(F3'A)$ | $-0.02 \pm 0.03$ | $1.7 \pm 0.3$ | $-0.00 \pm 0.01$ | 3-4 |
| $\alpha\beta\gamma(F20'A)$ | $0.01 \pm 0.00$ | $1.7 \pm 0.7$ | $-0.00 \pm 0.01$ | 4-5 |
| $\alpha\beta\gamma(F24'A)$ | $-0.0 \pm 0.0$ | $1.8 \pm 0.47$ | $-0.01 \pm 0.02$ | 7-8 |
| $\alpha\beta\gamma(Y25'A)$ | $0.04 \pm 0.05$ | $1.4 \pm 0.20$ | $-0.02 \pm 0.01$ | 3-5 |

Given that 0’ and 4’ mutations in mASIC1a alter channel gating, apparent pore diameter, and ion selectivity, and M1 aromatic residues are highly conserved (especially in ENaCs), we were surprised that 0’ and 4’ mutations in hENaC had no effect on ion selectivity (note that only the 4’ position in hENaC α is aromatic in nature, whereas 4’ is occupied by threonines in β and γ subunits). To further test the potential contributions of conserved 0’ and 4’ aromatic side chains to currents through hENaC, we measured single channel conductance in WT and mutant channels. We thus expressed hENaC WT, 0ʹ and 4ʹ mutants in *Xenopus laevis* oocytes and measured currents to obtain the single channel conductances of WT and 0ʹ/4ʹ mutants containing amino acid substitutions in one subunit (α) or two subunits (α and β). The single channel conductances of the 0ʹ mutants were significantly increased from 4.7 pS ± 1.0 in WT hENaC to 7.4 pS ± 3.2 in α(W0ʹF)βγ and 8.6 pS ± 1.0 in α(W0ʹF)β(W0ʹF)γ (Figure 2F/G and Table 4), whereas the single channel conductances of the 4’ single and double mutants were not significantly different from WT hENaC (Figure 3F/G and Table 4). These results implicate the highly conserved W0ʹ, but not the 4ʹ side chain in conductance.

Our data show that although hENaC M1 mutations can affect conductance, their impact on ion selectivity is generally small. We thus conclude that M1 contributes to ion selectivity to a lesser degree in hENaC than in mASIC1a.

### Determinants of ion selectivity in the pre-M1 segment of mASIC1a

We next wanted to expand our investigation into other regions possibly contributing to ion selectivity in ASICs and ENaCs. Because the most important determinants of ion selectivity in M1 were found to be located towards the intracellular end of M1, we wanted to probe side chains further upstream of these positions. Work by others had already highlighted the functional importance of this pre-M1 region [24, 25], and the latest cASIC structures reveal that residues in the N-terminus preceding M1 form part of the lower pore structure in ASICs [22]. We used comparative sequence analysis of 33 ASIC- and ENaC sequences, as well as other members of the DEG/ENaC superfamily (Figure 4A and Supplementary Figure 4), to identify potentially important determinants of ion selectivity in the pre-M1 segment. The resulting sequence logo indicates the degree of amino acid conservation in this region (Figure 4A).

**Figure 4:**
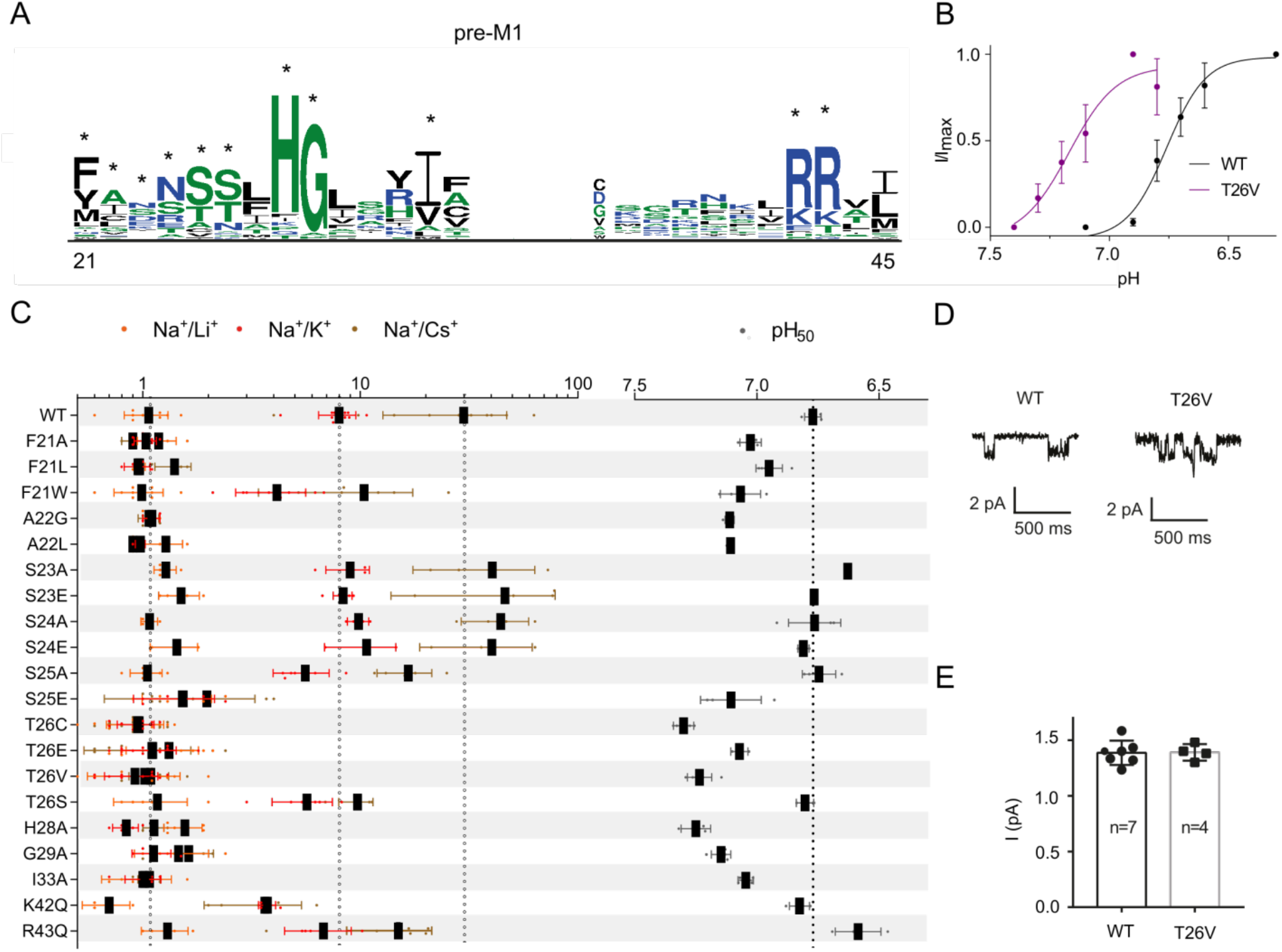
Numerous mASIC1a pre-M1 mutations have strong impact on ion selectivity and channel gating. (A) Sequence logo based on 33 ASIC, ENaC and related sequences (see Supplementary Figure 4). The height of each amino acid at a given position is proportional to its frequency at this position. The sequence logo was generated using the WebLogo 3 program (http://weblogo.threeplusone.com/create.cgi). The positions that were individually substituted in mASIC1a are highlighted with asterisks and positions are numbered based on the position in mASIC1a. (B) pH concentration response curves of WT mASIC1a and the T26V mutant. Data shown mean ± SD, n = 5-6. (C) Residues that occur frequently in the DEG/ENaC superfamily of ion channels were mutated in mASIC1a to the indicated amino acids to assess their contributions to ion selectivity and proton sensitivity. Relative ion permeability ratios (shown on a log scale for clarity) and half-maximal effective proton concentration values (pH_50_) of mASIC1a WT and pre-M1 mutants. Values are presented as mean ± SD, n ≥ 4. (D) Single channel traces recorded in *Xenopus laevis* oocytes expressing WT mASIC1a and the T26V mutant determined in outside-out patches. Cells were clamped at -60 mV and continuously perfused with pH 8 solution (in mM 140 NaCl, 3 MgCl_2_ and 5 HEPES and activated with pH 6. The intracellular solution contained (in mM) 110 NMDG, 2 MgCl_2_, 2 CaCl_2_, 10 HEPES, 5 EGTA, pH 7.3). (E) Averaged single channel current at -60 mV. Data shown as individual data points, with mean ± SD.

We started by investigating the N-terminal end of this sequence stretch, which in mASIC1a comprises a FASSST sequence motif: substitution of F21 (mASIC1a numbering) for the smaller L or A reduced relative ion permeability to unity and caused an alkaline shift in the proton sensitivity (pH_50_ = 6.8 ± 0.0 in WT, 7.0 ± 0.0 in F21A and 7.0 ± 0.1 in F21L), whereas substitution with the larger W decreased ion selectivity to a lesser degree, but caused a similar alkaline shift in pH_50_ (7.1 ± 0.1) (Figure 4C and Table 6). Substitution of the neighboring A22 for G and L also abolished ion selectivity and shifted the pH_50_ values to 7.1 ± 0.0 in A22G and 7.1 ± 0.0 in A22L. The S23A, S23E and S24A mutants were not significantly different from WT in terms of ion selectivity or the pH_50_ values, but the S24E mutant exhibited a modest increase in the Na^+^/K^+^ permeability ratio (10.7 ± 3.9). Previous work had suggested hASIC1a S25 to be phosphorylated [37], so we mutated S25 to A and E, in order to either remove a potential phosphorylation (A) or mimic the negative charge of a phosphate group (E). However, Na^+^ selectivity in S25A was only slightly decreased compared to WT (P_Na+/_K_+_ permeability = 5.6 ± 1.6, with WT-like proton sensitivity: pH_50_ = 6.8 ± 0.1), while the S25E mutant abolished Na^+^ selectivity and proton sensitivity was markedly increased (pH_50_ = 7.1 ± 0.1). This argues against a direct role for phosphorylation of S25 in channel function. By contrast, substitution of the neighboring T26 for C, V, or E reduced ion selectivity to unity and drastically shifted the pH_50_ to more alkaline pH (Figure 4B/C and Table 6), while the T26S ion selectivity profile resembled that of WT. For this mutant the pH_50_ value was not different from WT and it displayed only modest differences in the Na^+^/K^+^ permeability ratio (5.7 ± 1.8 compared to 6.8 ± 0.3 in WT) and in the Na^+^/Cs^+^ permeability ratio (9.7 ± 1.7 compared to 6.8 ± 0.3 in WT) (Figure 3C and Table 6). Together, this indicates that the T26 hydroxyl (or its T26S equivalent) are crucial to both Na^+^ selectivity and proton-dependent gating. However, we found that the isosteric T26V mutant did not alter the amplitude of Na^+^-mediated single channel currents, suggesting that the loss of selectivity in this mutant does not reduce relative permeability of Na^+^, but is rather more likely to increase relative permeability of K^+^ (Figure 4D/E).

**Table 6:** Relative ion permeabilities and pH_50_ values of mASIC1a mutants. Relative ion permeabilities and pH_50_ values of WT mASIC1a and mutants containing amino acid substitutions in the pre-M1 region. Data shown as mean ± SD with *n* ≥ 4, where *n* refers to the number of individual oocytes; n.d., not determined. Values for mutants were compared to WT using a one-way ANOVA with a Dunnett’s multiple comparisons test. *, *p* < 0.05; **, *p* < 0.01; ***, *p* < 0.001; ****, *p* < 0.0001.

| Construct | $P_{Na^+}/P_{K^+}$ | $P_{Na^+}/P_{Li^+}$ | $P_{Cs^+}/P_{Na^+}$ | pH<sub>50</sub> |
| --- | --- | --- | --- | --- |
| WT | 8.0 ± 1.6 | 1.1 ± 0.2 | 29.9 ± 17.2 | 6.8 ± 0.03 |
| F21A | 1.0 ± 0.1 **** | 1.2 ± 0.2 | 0.9 ± 0.1 ** | 7.0 ± 0.04 **** |
| F21L | 1.0 ± 0.1 **** | 1.0 ± 0.1 | 1.4 ± 0.3 ** | 7.1 ± 0.05 *** |
| F21W | 4.1 ± 1.5 **** | 1.0 ± 0.3 | 10.4 ± 7.0 ** | 7.1 ± 0.08 **** |
| A22G | 1.1 ± 0.1 **** | 1.1 ± 0.1 | 1.1 ± 0.1 ** | 7.1 ± 0.02 **** |
| A22L | 1.0 ± 0.1 **** | 1.3 ± 0.3 | 0.9 ± 0.0 *** | 7.1 ± 0.01 **** |
| S23A | 9.0 ± 2.0 | 1.3 ± 0.2 | 40.4 ± 22.9 | 6.6 ± 0.01 |
| S23E | 8.3 ± 0.8 | 1.5 ± 0.3 | 46.2 ± 32.4 | 6.8 ± 0.01 |
| S24A | 9.8 ± 1.1 | 1.1 ± 0.1 | 44.2 ± 15.1 | 6.8 ± 0.1 |
| S24E | 10.7 ± 3.9 ** | n.d. | 40.3 ± 21.5 | 6.8 ± 0.02 |
| S25A | 5.6 ± 1.6 ** | 1.1 ± 0.2 | 16.6 ± 4.7 | 6.8 ± 0.07 |
| S25E | 1.5 ± 0.6 **** | 1.5 ± 0.5 | 2.0 ± 1.3 **** | 7.1 ± 0.1 **** |
| T26C | 0.9 ± 0.2 **** | 1.0 ± 0.3 | 1.0 ± 0.2 **** | 7.1 ± 0.03 **** |
| T26E | 1.1 ± 0.3 **** | 1.3 ± 0.5 | 1.1 ± 0.6 **** | 7.2 ± 0.05 **** |
| T26V | 0.9 ± 0.3 **** | 1.0 ± 0.5 | 1.1 ± 0.2 **** | 7.2 ± 0.04 **** |
| T26S | 5.7 ± 1.8 ** | 1.2 ± 0.4 | 9.7 ± 1.7 * | 6.8 ± 0.04 |
| H28A | 0.8 ± 0.1 **** | 1.6 ± 0.3 | 1.1 ± 0.1 *** | 7.3 ± 0.06 **** |
| G29A | 1.1 ± 0.2 **** | 1.5 ± 0.6 | 1.6 ± 0.5 *** | 7.2 ± 0.04 **** |
| I33A | 1.0 ± 0.2 **** | 1.0 ± 0.4 | 1.1 ± 0.1 *** | 7.0 ± 0.03 **** |
| K42Q | 3.7 ± 0.3 **** | 0.7 ± 0.2 | 3.6 ± 1.7 *** | 6.8 ± 0.04 |
| R43Q | 6.8 ± 2.3 | 1.3 ± 0.3 | 15.0 ± 6.3 | 6.6 ± 0.09 *** |

The highly conserved H28 and G29 positions were recently shown to be located near the apex of a re-entrant loop formed by part of the cASIC1 N-terminus [22]. Consistent with the notion that both residues are crucial to function (and lining the permeation pathway), mutations at both sites disrupted ion selectivity and proton sensitivity: substitution of either of the two amino acids with A resulted in channels that were equally permeable to Na^+^, K^+^, Li^+^ and Cs^+^. Additionally, the pH_50_ values of both mutants were drastically left-shifted to 7.3 ± 0.1 for H28A and 7.2 ± 0.0 for G29A, respectively. This is in agreement with previous work on the equivalent positions in ENaC [38, 39]. While alanine substitution at the conserved I33 resulted in a similar phenotype (loss of selectivity, left-shifted pH_50_), mutation of K42 and R43 at the bottom of M1 showed a more modest reduction in Na^+^ selectivity and a less pronounced effect on proton dependence: the K42Q mutant showed WT-like gating with slightly reduced ion selectivity (P_Na+_/P_K+_ = 3.7 ± 0.3 and P_Na+_/P_Cs+_ = 3.6 ± 1.7), while the R43Q mutant was slightly less sensitive to protons (pH_50_ = 6.6 ± 0.1), and also displayed modestly reduced ion selectivity (P_Na+/_P_K+_ = 6.8 ± 2.3 and P_Na+/_P_Cs+_ = 15.0 ± 6.3). Interestingly, the mutants that exhibit large alkaline shifts in pH_50_ tend to also show distinct currents phenotypes, where the peak current was typically followed by a sustained current (Supplementary Figure 5). Together, this complements the latest structural findings, which suggest that these residues contribute to the ion permeation pathway (Supplementary Figure 5) and mutational disruptions appear to cause severe consequences, manifested in abolished ion selectivity and altered gating.

In summary, our data on mASIC1a clearly shows that the pre-M1 region is important for both ion selectivity and channel gating.

### Determinants of ion selectivity in the pre-M1 of hENaC

Intrigued by the pronounced effects of mutations in the stretch of amino acids comprising the FASSST sequence motif in the pre-M1 region of mASIC1a, we sought to probe the potential role of the equivalent amino acids in the hENaC α, β and γ subunits. To this end, alanine substitutions were introduced in one, two or all three subunits of the FASSST sequence equivalent in hENaC. Additionally, two threonines immediately following this stretch of amino acids in the β- and γ subunits of hENaC were mutated to valine (Figure 5A). None of the mutants were substantially different from WT hENaC in terms of the relative K^+^/Na^+^ and Cs^+^/Na^+^ current ratios, and Li^+^/Na^+^ current ratio was only modestly altered (Figure 5D). The only exception was the α(F62A)β(Y29A)γ double mutant, for which the Li^+^/Na^+^ current ratio was significantly increased from 1.2 ± 0.2 in WT to 3.7 ± 2.1 in α(F62A)β(Y29A)γ, *p* < 0.0001 (Figure 5D and Table 7). The α(F62A)β(Y29A)γ(Y32A) and α(T66V)β(T33V)γ(T36V) triple mutants did not generate detectable currents.

**Figure 5:**
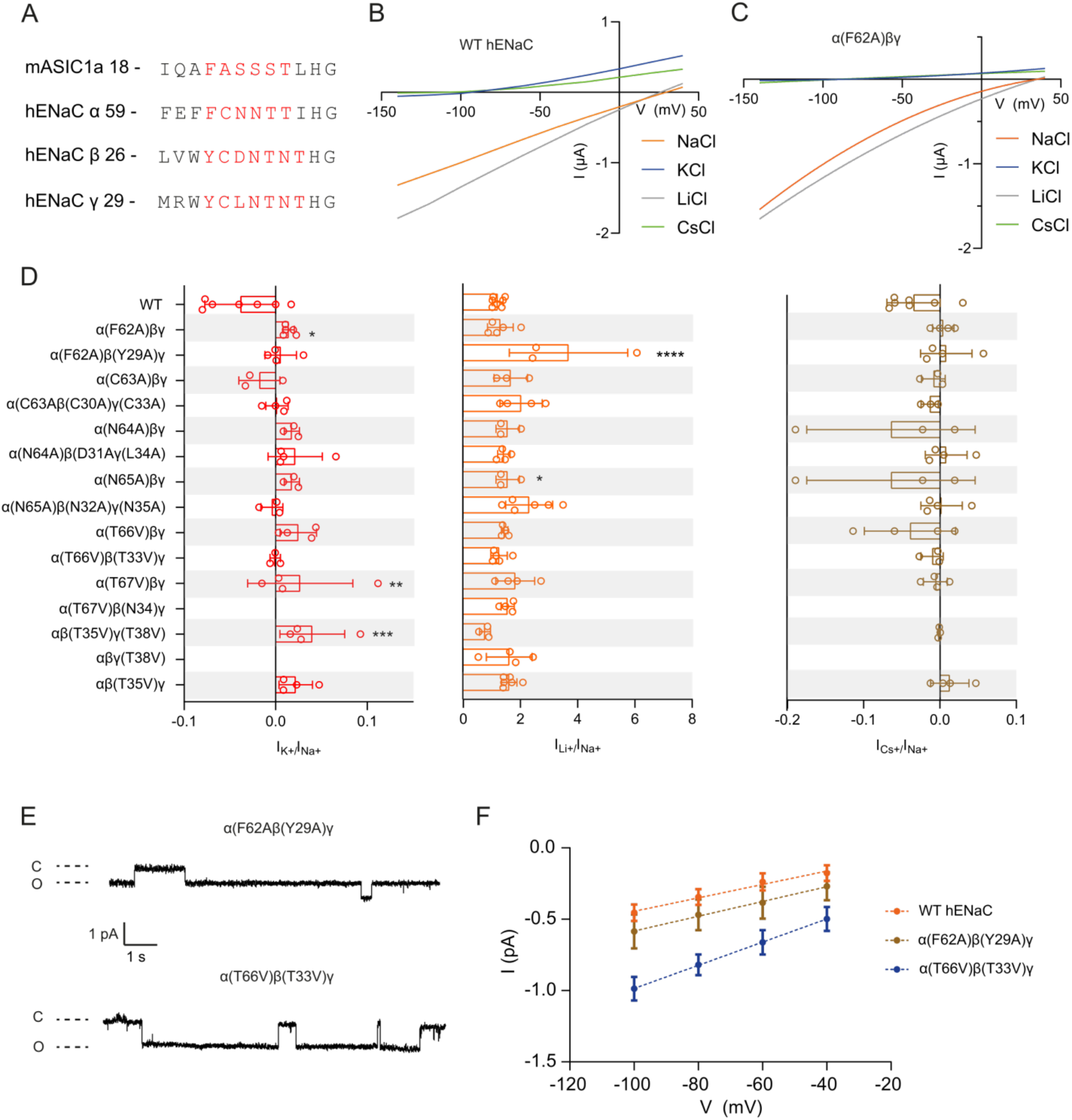
pre-M1 mutations in hENaC primarily alter conductance, but not ion selectivity. (A) Sequence alignment of part of the preM1 region of mASIC1a and hENaC. Residues that were mutated individually are highlighted in red. (B)(C) Current-voltages relationships of the amiloride-sensitive currents for WT hENaC (B) and the designated mutant (C). (D) Relative amiloride-sensitive I_K+_/_Na+_ (red), I_Li+_/I_Na+_ (orange) and I_Cs+_/I_Na+_ (brown) currents measured at –100 mV in WT hENaC and the designated mutants. Values shown as mean ± SD. Data were compared by a one-way ANOVA with a Dunnett’s multiple comparisons test for comparison with control (WT hENaC). *, *p* < 0.05; **, *p* < 0.01; ***, *p* < 0.001. (E) Single-channel currents of depicted hENaC mutants. Currents were recorded in *Xenopus laevis* oocytes expressing hENaC channels using the cell-attached patch-clamp configuration at a holding potential of +100 mV with 140 mM NaCl in the pipette. (F) Single-channel I/V relationships for the channels shown in (C). Data presented as mean ± SD. Slopes estimated by linear regression resulted conductances of: WT = 4.7 pS ± 1.0; α(T66V)β(T33V) = 8.6 pS ± 1.0; α(F62A)β(Y29A)γ = 5.2 pS ± 0.3. Data presented as mean ± SD for 4 – 7 patches. The value for α(T66V)β(T33V)γ is significantly greater than that of the WT channel (*p*-value = 0.0004).

**Table 7:** Relative amiloride-sensitive currents in WT hENaC and hENaC mutants. Relative amiloride-sensitive currents determined for WT hENaC and hENaC mutants containing the designated amino acid substitutions. Data shown as mean ± SD and *n* refers to the number of individual oocytes; n.d., not determined. Values for mutants were compared to WT hENaC using a one-way ANOVA with a Dunnett’s multiple comparisons test. *, *p* < 0.05; **, *p* < 0.01; ***, *p* < 0.001; ****, *p* < 0.0001, n = 3-7.

| Construct | $I_K/I_{Na}$<br>(-100mV) | $I_{Li}/I_{Na}$<br>(-100mV) | $I_{Cs}/I_{Na}$<br>(-100mV) |
| --- | --- | --- | --- |
| WT hENaC | $-0.04 \pm 0.04$ | $1.2 \pm 0.2$ | $-0.04 \pm 0.04$ |
| $\alpha\beta(T35V)\gamma$ | $0.02 \pm 0.02^*$ | $1.6 \pm 0.3$ | $0.01 \pm 0.03$ |
| $\alpha\beta\gamma(T38V)$ | n.d. | $1.6 \pm 0.8$ | n.d. |
| $\alpha\beta(T35V)\gamma(T38V)$ | $0.04 \pm 0.04^{***}$ | $0.8 \pm 0.2$ | $-0.00 \pm 0.00$ |
| $\alpha(T67V)\beta(N34A)\gamma$ | n.d. | $1.6 \pm 0.2$ | n.d. |
| $\alpha(T67V)\beta\gamma$ | $0.03 \pm 0.06^{**}$ | $1.8 \pm 0.7$ | $-0.01 \pm 0.02$ |
| $\alpha(T66V)\beta(T33V)\gamma$ | $-0.00 \pm 0.01$ | $1.3 \pm 0.3$ | $-0.01 \pm 0.02$ |
| $\alpha(T66V)\beta\gamma$ | $0.03 \pm 0.02^{**}$ | $1.5 \pm 0.1$ | $-0.04 \pm 0.06$ |
| $\alpha(N65A)\beta(N32A)\gamma(N35A)$ | $-0.00 \pm 0.01$ | $2.3 \pm 0.8^*$ | $0.00 \pm 0.03$ |
| $\alpha(N65A)\beta\gamma$ | $0.02 \pm 0.01$ | $1.6 \pm 0.4$ | $-0.06 \pm 0.11$ |
| $\alpha(N64A)\beta(D31A)\gamma(L34A)$ | $0.02 \pm 0.03^*$ | $1.4 \pm 0.2$ | $0.01 \pm 0.03$ |
| $\alpha(N64A)\beta\gamma$ | $0.02 \pm 0.01$ | $1.6 \pm 0.4$ | $-0.06 \pm 0.11$ |
| $\alpha(C63A)\beta(C30A)\gamma(C33A)$ | $0.00 \pm 0.01$ | $2.0 \pm 0.7$ | $-0.01 \pm 0.01$ |
| $\alpha(C63A)\beta\gamma$ | $-0.02 \pm 0.02$ | $1.7 \pm 0.6$ | $-0.01 \pm 0.02$ |
| $\alpha(F62A)\beta(Y29A)\gamma$ | $0.01 \pm 0.02$ | $3.7 \pm 2.1^{****}$ | $0.01 \pm 0.03$ |
| $\alpha(F62A)\beta\gamma$ | $0.01 \pm 0.01^*$ | $1.3 \pm 0.5$ | $0.00 \pm 0.01$ |

Although these data suggest that the pre-M1 segment does not make a major contribution to ion selectivity in hENaC, we next tested if the amino acids corresponding to the mASIC1a FASSST sequence contribute to ion conduction. We thus determined single channel conductances for i) α(F62A)βγ and α(F62A)β(Y29A)γ (Table 8), because the equivalent aromatic amino acids at this position in mASIC1a play a role in ion selectivity and proton sensitivity (Figure 4C); ii) mutants containing single T to V substitutions at all positions in this region containing the residue T (α(T66V)βγ, α(T66V)β(T33V)γ, α(T67V)βγ, αβ(T35V)γ and αβγ(T38V), Figure 5A). Single channel currents were not detectable for α(T67V)βγ, αβ(T35V)γ and αβγ(T38V), but there was a marked increase in the single channel conductance of the α(T66V)β(T33V)γ mutant compared to WT hENaC (Figure 5E/F and Table 8). By contrast, the α(F62A)β(Y29A)γ double mutant showed only a small, but insignificant increase in single channel amplitude (Figure E/F and Table 8).

**Table 8:** Single channel conductances of WT hENaC and pre-M1 mutants. WT and mutant hENaC RNA was injected in *Xenopus laevis* oocytes and single-channel conductances were determined in the cell-attached patch clamp configuration with 140 mM extracellular Na^+^. Single channel conductances presented as mean ± SD and were compared to WT ENaC using a one-way ANOVA with a Dunnett’s multiple comparisons test. *, *p* < 0.05; **.

| Construct | Single channel conductance (pS) | *n* |
| --- | --- | --- |
| WT hENaC | 4.7 ± 1.0 | 6 |
| $\alpha$(F62A)$\beta\gamma$ | 4.5 ± 1.0 | 5 |
| $\alpha$(F62A)$\beta$(Y29A)$\gamma$ | 5.2 ± 0.3 | 5 |
| $\alpha$(T66V)$\beta\gamma$ | 7.1 ± 0.6 * | 7 |
| $\alpha$(T66V)$\beta$(T33V)$\gamma$ | 8.1 ± 0.6 ** | 5 |

Together, our data suggest that the pre-M1 region in hENaC is not a major determinant of ion selectivity but contributes to ion conduction.

## Discussion

Recent advances in structural biology have resulted in the publication of multiple structures of chick ASIC1 and hENaC [15, 16, 21, 22, 40, 41]. However, structures of open, full-length ASICs have not been obtained and especially the conformation of the lower pore varies considerably among ASIC structures [15, 21, 22, 41]. Similarly, the resolution of the pore-forming M1 and M2 segments in the available ENaC structures is comparatively low, giving rise to uncertainty regarding their absolute and relative positions, also with respect to the (unresolved) N- and C-termini [16, 40]. It is therefore difficult to reconcile the stark differences in ion selectivity among these channel types solely based on available structural data. Here we use a combination of electrophysiological approaches to assess the contribution of M1 and pre-M1 to ion selectivity in mASIC1a and hENaC. We demonstrate that these regions contribute differently in both channel types: while mutations have severe consequences for ion selectivity in mASIC1a, they mostly affect conductance in hENaC. These findings support the notion that despite their similarities in sequence and structure, ASICs and ENaCs achieve ion selectivity via different mechanisms.

### Both M1 and pre-M1 segments of mASIC1a contain key determinants of ion selectivity

In agreement with previous work, we find that mutations [24, 42] at the three native cysteines in ASIC1a M1 (C3ʹL, C13ʹW and C15ʹF) do not have a major impact on ion selectivity or channel gating (Figure 2C). Similarly, other mutations along M1 between F23ʹ and G6ʹ do not change ion selectivity. By contrast, we find that mutations of aromatic side chains towards the intracellular end of M1 (F4ʹ and W0ʹ) abolish sodium selectivity. We also observe significant changes in apparent proton sensitivity with W0ʹ mutants, which is consistent with earlier reports and supportive of the notion that this highly conserved side chain is important for channel gating [43, 44]. F4ʹ mutants, on the other hand, had WT-like gating and only affected ion selectivity. We therefore considered the F4ʹA mutant a reasonable model to test the notion that reduced bulk in the lower M1 may lower ion selectivity by increasing pore diameter. The finding that larger cations, such as MA^+^, EMA^+^ and EA^+^ permeate F4ʹA mutant channels more readily than WT channels lends direct support to this hypothesis. We conclude that the two aromatic residues in lower M1, W0ʹ and F4ʹ, indirectly contribute to the pore diameter, and likely overall structure of the mASIC1a lower pore.

Additionally, these results pointed towards the lower pore as a major determinant for ion selectivity in ASICs. Although the exact location of the selectivity filter in ASICs has remained a matter of debate, these findings support the notion that the selectivity filter might be located in the lower pore [17, 23]. This led us to postulate that M2-E18ʹ and M2-D21ʹ are likely to form the selectivity filter in ASICs. However, the recent cASIC1 structures clearly show that at least in closed and desensitized conformations, it is the pre-M1 region that lines the lower ion permeation pathway, thus calling into question whether M2-E18ʹ and M2-D21ʹ make direct contributions to ion selectivity in ASICs [22]. The structures raise the possibility that the ASIC selectivity filter might be, at least in part, formed by the pre-M1 segment, specifically the narrow constriction formed at the histidine from the conserved HG motif that is involved in a network of both intra- and inter-subunit interactions [22]. This is in agreement with our finding that the pre-M1 segment of mASIC1a is extremely sensitive to mutations, such that even minor mutational disruptions abolish ion selectivity, including those to the HG motif (Figure 4C). In fact, this protein segment contains numerous side chains that, when mutated, have drastic consequences for both ion selectivity and channel gating, therefore underlining its functional importance. By comparison, we found relatively fewer side chains in the pore-lining M2 segment to affect ion selectivity [17, 23].

Together, these data are in agreement with the possibility that the selectivity filter is located at the lower end of the pore, likely formed by the pre-M1 segment and at least indirectly supported by E18ʹ and D21ʹ in M2 and W0ʹ and F4ʹ in M1. However, in the absence of an open channel structure with a fully resolved pre-M1 region we cannot rule out the possibility that other side chains, including M2-E18ʹ and M2-D21ʹ, may directly line the permeation pathway in the channel open state. Interestingly, none of the mutations tested in this study increases ion selectivity of ASIC1a. As ENaC is more Na+ selective than ASIC, but none of the numerous ASIC mutations, even to ENaC-like identity, markedly increase Na+ selectively, ASICs seem to have developed – independently from ENaCs – a fragility in their molecular blueprint for ion selectivity.

### ENaC ion selectivity is comparatively impervious to mutations in M1 and pre-M1

Although structural information on the ENaC pore is of relatively low resolution [16, 40], there is strong functional evidence that the G/SXS motif in the middle of the pore is critical for ion selectivity in ENaCs [29, 36, 38, 45]. Additionally, and in contrast to the findings for ASICs, the negatively charged side chains in the M2-18ʹ and M2-21ʹ positions do not appear to play a crucial role in ion selectivity [29]. Yet it is interesting to note that mutations to W0ʹ in the ENaC *α*and *β* subunits affected both selectivity and single channel conductance. Although the triple mutant was non-functional, this points towards W0ʹ being - directly or indirectly - involved in ion conduction in ENaCs (Figure 2), although likely to a lesser degree than in ASICs. Furthermore, important functional contributions by side chains in the (putative) lower pore of ENaCs are highlighted by the finding that mutations to the conserved HG motif affect gating by decreasing ENaC open probability [38] and mutations of the G in this motif causing pseudohypoaldosteronism type 1 (PHA1) in humans [39]. Our results expand on these results by showing that selected pre-M1 mutations can affect ion selectivity or conductance (Figure 5). This shows that in contrast to ASICs, mutations at a few positions in both the middle and the lower part of the pore have relatively minor effects on ion selectivity but contribute significantly to conduction in ENaCs. Additionally, reducing the bulk of aromatic side chains along the entire length of at least the *α* subunit M1 in hENaC, such as W0ʹ, F8ʹ, F19ʹ, Y22ʹ and Y23ʹ can lead to (albeit small) changes in ion selectivity.

This indicates that although the overall ion selectivity profile is maintained in most ENaC mutants, subtle changes can be elicited by mutations throughout the M1 and pre-M1 segments. This is in contrast to ASICs, where mutations in the upper and middle sections of both M1 and M2 do not result in changed ion selectivity (the only exception being C15ʹF, see Figure 2C). We thus conclude that the open pore structures of ASICs and ENaCs likely display important structural differences.

### Limitations and outlook

In the majority of our work on ASICs, we rely on relative ion permeability ratios as a proxy for ion selectivity. More subtle changes in conductance, as revealed for selected ENaC mutants, are therefore harder to assess in measurements of relative ion permeability ratios. But previous studies demonstrate that relative ion permeability measurements can result in robust estimates of ion selectivity [17, 19, 23, 36]. Also, and in line with work by others [29], some of the ENaC triple mutants did not generate measurable currents. We therefore cannot exclude the possibility that we may underestimate the contributions of some side chains to selectivity and/or conductance. Additionally, it has been reported that the ENaC pore helices might be arranged asymmetrically and that mutations at equivalent sites may not cause equivalent effects [46]. Finally, the lack of definitive structural information on the open pore structures of both ASICs and ENaCs means that it is difficult to interpret our functional data in detailed structural context.

Overall, however, the data at hand clearly support the notion of ASICs and ENaCs achieve sodium selectivity by different mechanisms. Our work therefore highlights how related channels can achieve different functional properties despite obvious sequence conservation in e.g. the (at least partially) pore-lining M2 segments. Instead, the data indicate that the non-pore-lining M1 segment indirectly contributes to ion selectivity in both ASICs and ENaCs.

## Acknowledgements

This work was supported by Lundbeckfonden (R171-2014-558, T. Lynagh; R139-2012-12390, S.A. Pless), the Danish Council for Independent Research (4092-00348B, T. Lynagh) and Carlsbergfondet (2013_01_0439, S.A. Pless).

## Author contributions

Z.P. Sheikh, T. Lynagh, W. Wulf and S. Friis conducted functional experiments; Z.P. Sheikh, T. Lynagh, W. Wulf, S. Friis and M. Althaus analyzed data; Z.P. Sheikh, T. Lynagh, S. Friis, M. Althaus and S.A. Pless conceptualized the work. Z.P. Sheikh and S.A. Pless drafted the manuscript and finalized it with input from all authors.

