## Supplementary information for "The M1 and pre-M1 segments contribute differently to ion selectivity in ASICs and ENaCs"

### Differential contributions of M1 and pre-M1 to ion selectivity in ASICs and ENaCs

Contents:

- Figure S1
- Figure S2
- Figure S3
- Figure S4
- Figure S5
- Figure S6
- Figure S7

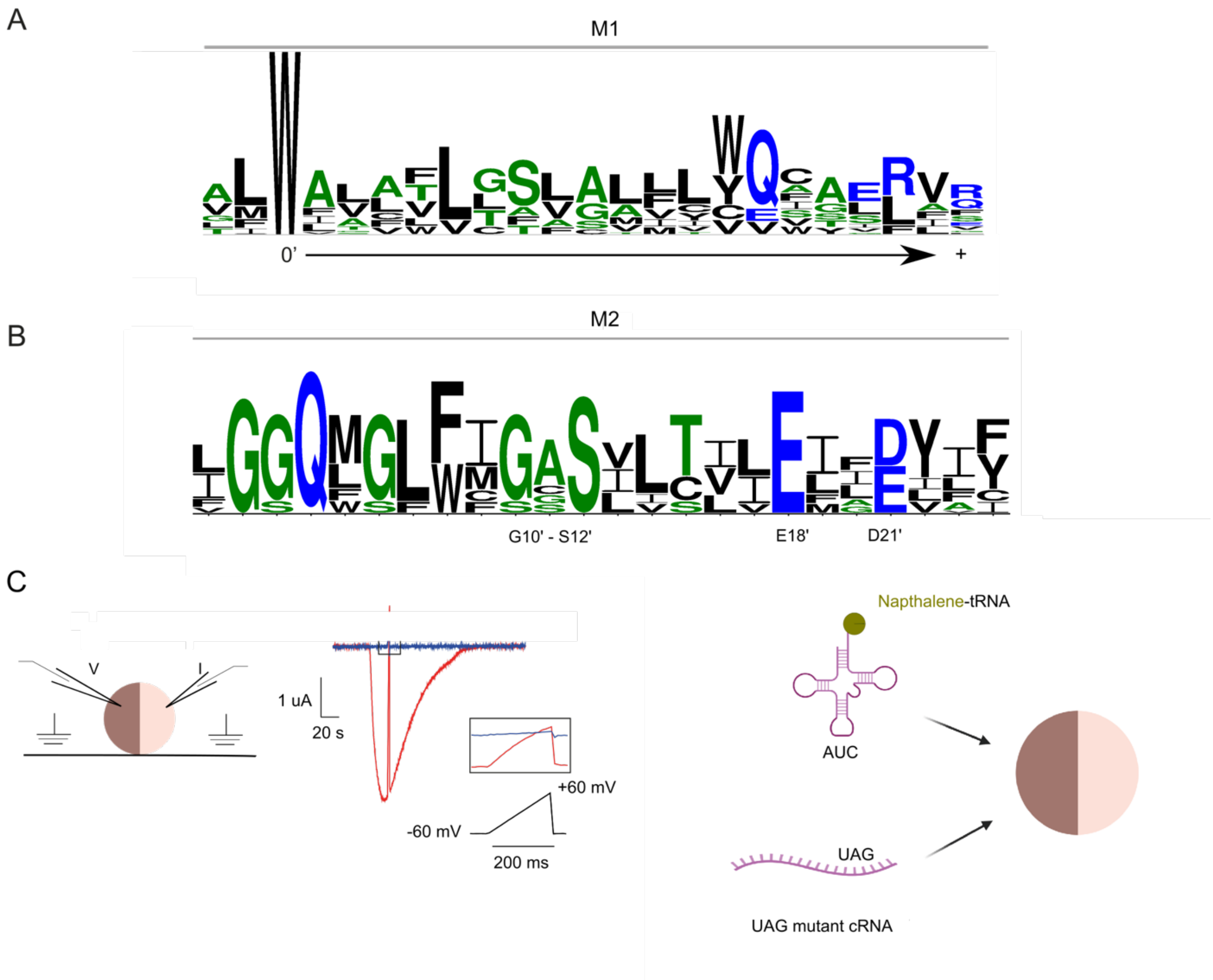


**Supplementary Figure 1: M1 and M2 sequence logos and overview of nonsense suppression approach.** (A) Sequence logo of M1 and M2 based on mammalian ASIC an ENaC sequences (see Supplementary Figures 6 and 7 below). Residues are numbered according to position in mASIC1a. The height of each residue is proportional to its frequency at this position. Here, we have employed a numbering system for M1 in which equivalent residues from ENaCs and ASICs are referred to with the same number. The conserved W in M1 is designated the 0’ position. (B) Experimental approach used to determine ion selectivity of mASIC1a constructs. cRNA encoding WT or mutant mASIC1a was injected into *Xenopus laevis* oocytes and reversal potentials with extracellular $Na^+$, $K^+$, $Li^+$, and $Cs^+$ were determined using a 200 ms voltage ramp from -60 to + 60 mV during the peak current. Currents during the voltage ramps at pH 7.4 (blue) were subtracted from currents during activating pH (red). (C) Incorporation of the non-canonical amino acid, naphthalene, was achieved via the nonsense suppression method in *Xenopus laevis* oocytes. The suppressor tRNA, THG73, lacking its terminal CA dinucleotide was enzymatically ligated to the ncAA naphthalene attached to a CA dinucleotide using an RNA ligase. The tRNA carrying naphthalene was injected into *Xenopus laevis* oocytes together with mASIC1a mRNA containing an amber stop codon at the site of interest.

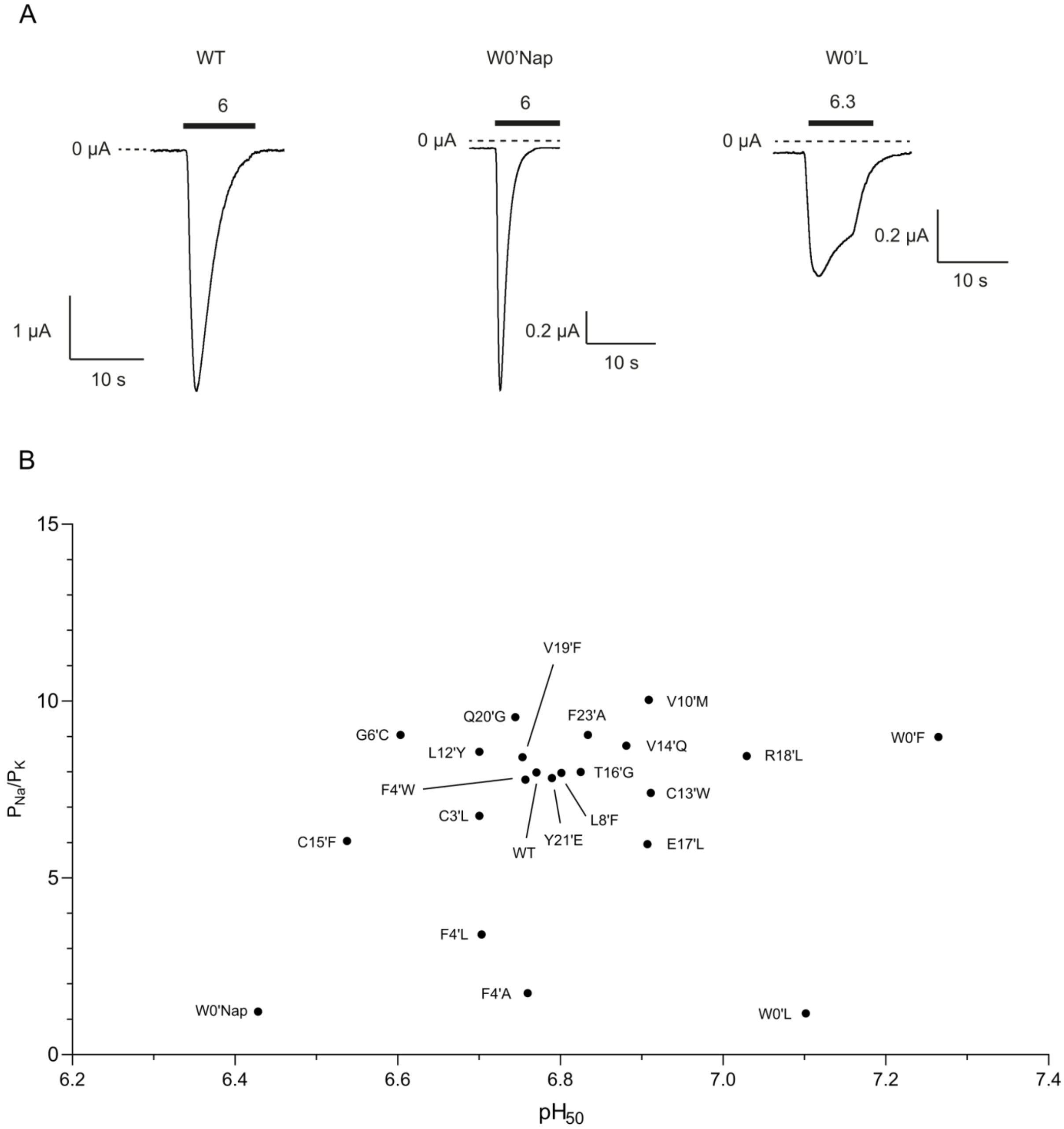


**Supplementary Figure 2: Current traces of WT and designated mutant mASIC1a channels and effects of M1 mutations on ion selectivity and $pH_{50}$.** (A) mRNA encoding WT or mutant mASIC1a channels were injected into *Xenopus laevis* oocytes, and currents were recorded using two-electrode voltage clamp. Cells were clamped at -20 mV and continuously perfused with ND96 solution (pH 7.4) and currents were elicited upon switching to ND96 solution with the designated pH. (B) Effects of M1 mutations on relative $Na^+$ over $K^+$ permeability ratios and $pH_{50}$ summarized with a scatter plot.

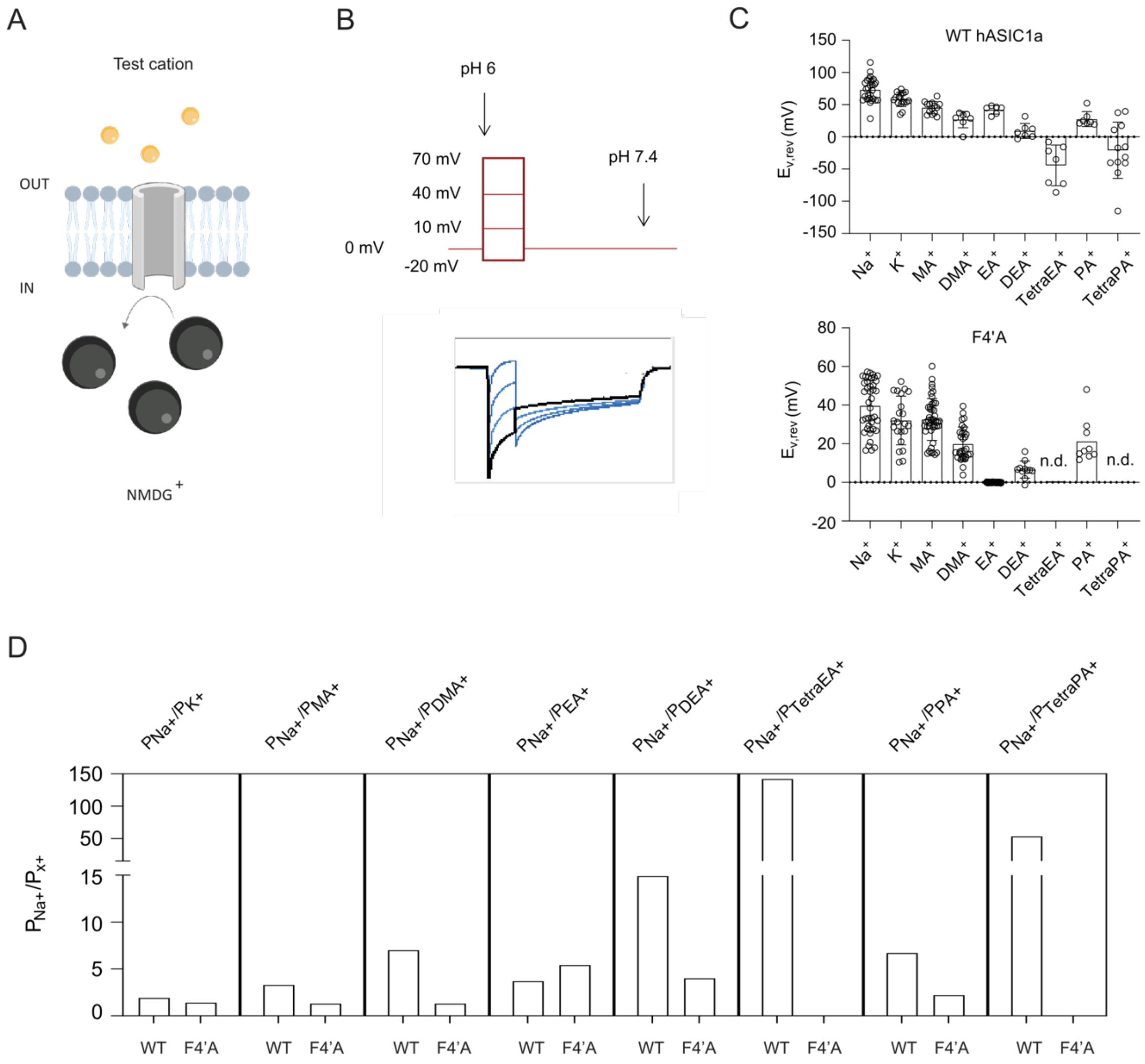


**Supplementary Figure 3: Automated patch clamp recordings using the SynchroPatch 384PE.** (A) Schematic illustration of the experimental setup. The intracellular solution contained $NMDG^+$ as the only cation and was selected based on its large size. The external side contained an ECS with one monovalent test cation. (B) The voltage protocol used to determine reversal potentials. ASIC1a K/O HEK-293T expressing either WT hASIC1a or the hASIC1a F50A mutant were clamped at 0 mV and currents were recorded by stepping to the four voltages, -20, 10, 40 and 70 mV, in four sweeps upon the simultaneous application of a saturating concentration of protons (pH 6). Screen shot of an example current trace carried by extracellular $Na^+$ is shown in the inset below the protocol. (C) Reversal potentials estimated with different extracellular monovalent cations using the protocol shown in (B). Abbreviations: $MA^+$: methylamine+, $DMA^+$: dimethylamine+, $EA^+$: ethylamine+, $DEA^+$: diethylamine+, $TEA^+$: trieethylamine+, $TetraEA^+$: tetraethylamine+, $PA^+$: propylamine+, $DPA^+$: dipropylamine+, and $TetraPA^+$: tetrapropylamine+. (D) Relative ion permeability ratios estimated for WT hASIC1a and the F4'A mutant, respectively.

```
Crassostrea_EKC42481.1     MLNNTSLHGVSKIS-----RGKTICHGVFWAVMTSVMLALLITVFVLYSLDYFLYETLI
Crassostrea_EKC39945.1     FADKTAMQGVGYIS-----SAKYWYSRAIWVFLLLVAMGWMVFHLYYLISQFTDLPVQT
Crassostrea_EKC42329.1     WLSAVDIHGLGIAG-----TTTHVLETSIWIIISCVAFSGFVFLVYLNFSVYYLQSSSF
XP_023240338.1             YEKRSNTKPPYIIA----WENSSKNKKK--LSRKKCSLSGFTYSSYKFLTNFFEYPVVV
Crassostrea_17580-23158    YGNSASFHGLRFVT----DPLANKPRRLIWLCLLTACLAVLVYQIVDRVTHFYSYPVTV
XP_023226921.1             FLGSSSVIGLSQIT-----KSRSIVRKLLWLAVLVTGLTFCAIESHKFMREFYKYPVVV
ENaCg_HUMAN                YCLNTNTHGCRRIV-----VSRGRLRRLLWIGFTLTAVALILWQCALLVFSFYTVSVSI
ENaCb_HUMAN                YCDNTNTHGPKRII----CEGPK--KKAMWFLLTLLFAALVCWQWGIFIRTYLSWEVSV
ENaCa_HUMAN                FCNNTTIHGAIRLV----CSQHNRMKTAFWAVLWLCTFGMMYWQFGLLFGEYFSYPVSL
ENaCd_HUMAN                FCTNATIHGAIRLV----CSRGNRLKTTSWGLLSLGALVALCWQLGLLFERHWHRPVLM
Crassostrea_EKC33938.1     FINETSFTAIARIV-----KANTILKKIIWLILVMAMMAWLTIQCYWLLDKYFSYPVEV
HyNaC2_157886734           YVQTSTLHGFRFIF-----MDTFIVRRVLWTILTLTMATIFFKELRNSINLFYEYPFTT
HyNaC5_289169249           YIEASTLHGYAYVC-----SDTFYIRRVLWAVLMLLGGIYFVFKLKVGIIEYFQYPFST
HyNaC6_594592304           YVESSTLHGFCYVC-----GDTFLVRRVLWALLMILGAIYFIIKLRYGIEEYLNYPFST
HyNaC7_594592306           YIESSTLHGFCYVC-----MDTFLGRRLIWAVLMILGAIYFIFKLRYGIKEYFDYPFST
ASIC2b_HUMAN               SLSRAKLHGLRHMCAGRTAAGGSFQRRALWVLAFCTSFGLLLSWSSNRLLYWLSFPSHT
bASIC_HUMAN                FAISTSFHGIHNIV-----QNRSKIRRVLWLVVVLGSVSLVTWQIYIRLLNYFTWPTTT
Crassostrea_EKC39946.1     NSRNSGLFGLGS---------------SPWEA---------------------------
ASIC1a_HUMAN               FASSSTLHGLAHIF----SYERLSLKRALWALCFLGSLAVLLCVCTERVQYYFHYHHVT
ASIC1b_HUMAN               FANSCTLHGTNHIF----VEGGPGPRQVLWAVAFVLALGAFLCQVGDRVAYYLSYPHVT
ASIC2a_HUMAN               FANTSTLHGIRHIF----VYGPLTIRRVLWAVAFVGSLGLLLVESSERVSYYFSYQHVT
ASIC3_HUMAN                FASNCSMHGLGHVF----GPGSLSLRRGMWAAAVVLSVATFLYQVAERVRYYREFHHQT
ASIC4_HUMAN                FASTSTLHGLGRAC----GPGPHGLRRTLWALALLTSLAAFLYQAAGLARGYLTRPHLV
Crassostrea_EKC22488.1     LGSESNAHGLAKIA-----MSRKTKRKVMWSLLVIIGFTAAAIHLSFLVIKYLQYNVVE
XP_023219716.1             IFKESSIPGVNNIA-----AAKGKFERFIWIFIVICCLTGFIYQVTLFLRHYYNYPTRV
XP_023215424.1             FAYRSSAHGIPRIA-----SSQNRFRRFMWIIVFLVAVAGFAYHSIYLILTYLSYPRMT
XP_023244170.1             FAYRSSAHGVQRIA-----SSQDNARRLMWSVVFLFAIAGCGFHSVYLILTYLSYPRMT
HyNaC8_594592308           MAVNSSFHGINYIC-----DSTYKVRRIIWIVVTLTAMLYAMREVYESTRKYLNYPVST
HyNac9_594592310           LIKNVSFHGLSYVA----DKRNNYFRRAIWFLITVGAFIYAVEKVYESTVNYFSYPFKT
HyNaC10_594592312          AGENISIHGLSHVF----DKRENFICRTVWLLITIAAFGYAVQKVYESTLNYFSYPFST
HyNaC4_157886739           MIDNSSFHGISYIA----GKENHFIRRTIWLLITMTAFGYAAQKVYESTVNYFSFPIST
HyNaC3_157886737           MIDNSSFHGLSYIF-----DKRHSIRRTIWFFITIAAFVYAMQKVYESTMNYFSYPFYT
HyNaC11_594592314          MIDNSSFHGLSYIF-----DKRHSVRRTIWFFITIAAFAYAMQKVYESTMNYFSYPFYT
```

**Supplementary Figure 4: Sequences used for the generation of the sequence logo shown in Figure 4.**

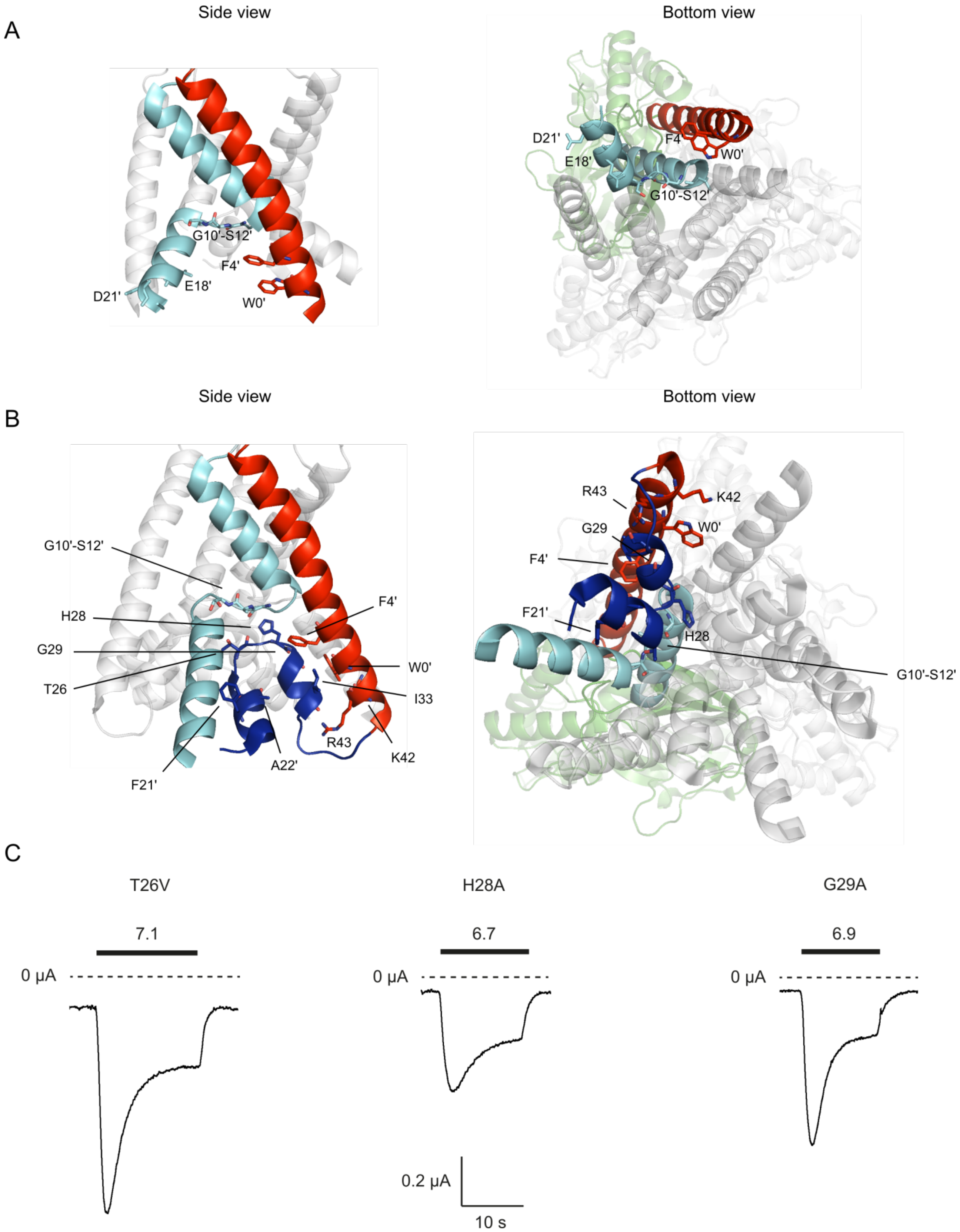


**Supplementary Figure 5: cASIC1 structures and example current traces of selected pre-M1 mASIC1a mutants.** Side view (left panel) and bottom view (right panel) of cASIC1 structures in an open (PDB: 4NTW) (A) and desensitized state (PDB: 6VTK) (B) with residues critical for ion selectivity shown as sticks. Blue: N, red: O. (C) Example traces of designated mASIC1a pre-M1 mutants recorded with two-electrode voltage clamp at -20 mV in *Xenopus laevis* oocytes (applied pH defined above the black bar indicating length of ligand application).

| | |
|---|---|
| ASIC1a_HUMAN | ALWALCFLGSLAVLLCVCTERVQ |
| ASIC2a_HUMAN | VLWAVAFVGSLGLLLVESSERVS |
| ASIC3_HUMAN | GMWAAAVVLSVATFLYQVAERVR |
| ASIC4_HUMAN | TLWALALLTSLAAFLYQAAGLAR |
| ASIC5_HUMAN | VLWLVVVLGSVSLVTWQIYIRLL |
| ASIC1a_MOUSE | ALWALCFLGSLAVLLCVCTERVQ |
| ASIC2a_MOUSE | VLWAVAFVGSLGLLLVESSERVS |
| ASIC3_MOUSE | GLWATAVLLSLAAFLYQVAERVR |
| ASIC4_MOUSE | TLWALALLTSLAAFLYQAASLAR |
| ASIC5_MOUSE | VIWLAVVLGSVSLLVWQIYSRLV |
| ASIC1a_RAT | ALWALCFLGSLAVLLCVCTERVQ |
| ASIC2a_RAT | VLWAVAFVGSLGLLLVESSERVS |
| ASIC3_RAT | GLWATAVLLSLAAFLYQVAERVR |
| ASIC4_RAT | TLWVLALLTSLAAFLYQAASLAR |
| ASIC5_RAT | VIWLSVVLGSVSLLVWQIYSRLV |
| ENaCα_HUMAN | AFWAVLWLCTFGMMYWQFGLLFG |
| ENaCβ_HUMAN | AMWFLLTLLFAALVCWQWGIFIR |
| ENaCγ_HUMAN | LLWIGFTLTAVALILWQCALLVF |
| ENaCα_MOUSE | AFWAVLWLCTFGMMYWQFALLFE |
| ENaCβ_MOUSE | AMWFLLTLLFACLVCWQWGVFIQ |
| ENaCγ_MOUSE | LLWIAFTLTAVALIIWQCALLVF |
| ENaCα_RAT | AFWAVLWLCTFGMMYWQFALLFE |
| ENaCβ_RAT | AMWFLLTLLFACLVCWQWGVFIQ |
| ENaCγ_RAT | LLWIAFTLTAVALIIWQCALLVF |

**Supplementary Figure 6: Sequences used for the generation of the sequence logo of M1 segment shown in Supplementary Figure 1.**

| | |
|---|---|
| ASIC1a_HUMAN | IGGQMGLFIGASILTVLELFDYAY |
| ASIC2a_HUMAN | IGGQMGLFIGASILTILELFDYIY |
| ASIC3_HUMAN | IGGQMGLFIGASLLTILEILDYLC |
| ASIC4_HUMAN | LGGQMGLFIGASILTLLEILDYIY |
| ASIC5_HUMAN | LGGQLGLFCGASLITIIEIIEYLF |
| ASIC1a_MOUSE | IGGQMGLFIGASILTVLELFDYAY |
| ASIC2a_MOUSE | IGGQMGLFIGASILTILELFDYIY |
| ASIC3_MOUSE | IGGQMGLFIGASLLTILEILDYLC |
| ASIC4_MOUSE | LGGQMGLFIGASILTLLEILDYIY |
| ASIC5_MOUSE | VGGQLGLFCGASLITIIEIIEYFF |
| ASIC1a_RAT | IGGQMGLFIGASILTVLELFDYAY |
| ASIC2a_RAT | IGGQMGLFIGASLLTILELFDYIY |
| ASIC3_RAT | IGGQMGLFIGASLLTILEILDYLC |
| ASIC4_RAT | LGGQMGLFIGASILTLLEILDYIY |
| ASIC5_RAT | VGGQLGLFCGASLITIIEIIEYLF |
| ENaCα_HUMAN | LGSQWSLWFGSSVLSVVEMAELVF |
| ENaCβ_HUMAN | LGGQFGFWMGGSVLCLIEFGEIII |
| ENaCγ_HUMAN | FGGQLGLWMSCSVVCVIEIIEVFF |
| ENaCα_MOUSE | LGSQWSLWFGSSVLSVVEMAELIF |
| ENaCβ_MOUSE | LGGQFGFWMGGSVLCLIEFGEIII |
| ENaCγ_MOUSE | FGGQLGLWMSCSVVCVIEIIEVFF |
| ENaCα_RAT | LGSQWSLWFGSSVLSVVEMAELIF |
| ENaCβ_RAT | LGGQFGFWMGGSVLCLIEFGEIII |
| ENaCγ_RAT | FGGQLGLWMSCSVVCVIEIIEVFF |

**Supplementary Figure 7: Sequences used for the generation of the sequence logo of M2 shown in Supplementary Figure 1.**
